# A pooled screening approach reveals bacterial chemoreceptors for short-chain carboxylic acids

**DOI:** 10.64898/2026.04.10.717710

**Authors:** Hiroya Oka, Dung Hoang Anh Mai, Sora Yu, Nicolas Grosjean, Zong-Yen Wu, Nam Ngoc Pham, David Robinson, Yasuo Yoshikuni, Nigel Mouncey, Tomoya Honda

**Affiliations:** US Department of Energy Joint Genome Institute, Lawrence Berkeley National Laboratory, Berkeley, CA 94720; Environmental Genomics and Systems Biology Division, Lawrence Berkeley National Laboratory, Berkeley, CA 94720; Biological Systems and Engineering Division, Lawrence Berkeley National Laboratory, Berkeley, CA 94720, USA; US Department of Energy Center for Bioenergy and Bioproducts Innovation (CABBI), Lawrence Berkeley National Laboratory, Berkeley, CA 94720, USA; Energy and Biosciences Institute, University of California, Berkeley, Berkeley, CA 94720; Global Institution for Collaborative Research and Education, Hokkaido University, Hokkaido 060-8589, Japan

## Abstract

Bacterial chemotaxis is a key process in host colonization and virulence, mediated by large repertoires of chemoreceptors. Despite their physiological and ecological importance, mapping these chemoreceptors to their cognate metabolic ligands remains a major challenge due to the vast number of potential interactions. To address this, we developed a pooled screening assay that enriches functional chemoreceptors from a gene library. Using this approach, we identified a previously uncharacterized group of chemoreceptors in *Pseudomonas* species with Cache_3–Cache_2 domains that sense short-chain C3 carboxylic acids. Sequence and computational structural analyses revealed that these chemoreceptors exhibit domain features similar to a recently reported C1 formate chemoreceptor despite substantial sequence divergence. Functional assays of representative chemoreceptors confirmed robust chemotactic responses to C3 carboxylic acids, with limited responses to formate. Integrating structural and molecular dynamics analyses suggests that increased binding pocket size and altered flexibility, relative to the formate chemoreceptor, facilitate recognition of larger C3 carboxylic acid ligands. Together, our approach provides a simple and scalable framework for mapping ligand–chemoreceptor interactions and enables systematic characterization across diverse metabolites.

## Introduction

Bacteria must sense and respond to external stimuli to grow and survive. These processes shape cellular physiology and interactions with hosts, including biofilm formation and virulence^1–3^. Over the past several decades, genetic and biochemical studies have uncovered diverse signal transduction systems in bacteria^4,5^. However, a major challenge remains in mapping metabolic ligands to their corresponding sensor proteins across a large number of possible interactions^6^. While AI-based approaches are advancing rapidly, experimental characterization remains essential for defining the functions of sensor proteins and improving predictive models.

Chemotaxis is a key sensory system that couples environmental sensing with bacterial behavior^7^. Through chemotaxis, cells detect chemical gradients and navigate toward favorable niches or away from adverse niches. This system is widely conserved among bacteria and provides competitive advantages by serving as an initial stage of colonization and virulence^8–13^. Although the molecular mechanisms of chemotaxis have been extensively studied in model organisms, these systems are highly diversified across bacterial species^14^. One of the major differences is the abundance and diversity of chemoreceptors, which determine the range of molecules that bacteria can detect and respond to. For example, rhizobacteria possess large repertoires of chemoreceptor genes, with an estimate of 23 genes per genome^15^. Agriculturally relevant bacteria, including plant pathogens and nitrogen-fixing species, harbor as many as ∼50 chemoreceptors^15,16^. Similarly, many human-associated and pathogenic bacteria encode a number of chemoreceptors^3^; for example, *Pseudomonas aeruginosa* encodes 26 chemoreceptors^17^. This is in contrast to well-studied species such as *Escherichia coli, Salmonella*, and *Bacillus subtilis* that have fewer than 10 chemoreceptors^14^. Understanding these diverse chemosensory systems is necessary to explain bacterial ecology and host associations.

Despite their physiological and ecological importance, the functions of most chemoreceptors in bacteria remain unknown. There are several factors that contribute to this limitation. First, the absence of genetic engineering tools makes it challenging to generate chemoreceptor mutants in non-model bacteria. Second, even if a mutant is created, functional redundancy among multiple chemoreceptors complicates phenotypic interpretation. Third, while in vitro assays^18,19^ and *E. coli*-based chimeric receptor constructions^20–24^ have provided valuable high-throughput approaches for ligand identification, their application can be limited by challenges associated with protein expression, folding, and purification. Last, although AI-based prediction holds great promise, their predictive power remains limited for chemoreceptors due to their complexity as small sequence variations can alter ligand specificity, while distantly related chemoreceptors can recognize similar ligands^14,25,23^.

To address these challenges, we developed a new high-throughput screening workflow to systematically characterize chemoreceptor functions. We generated an engineered bacterial population harboring a chemoreceptor gene library and screened for functional ones through enrichment assays. As a proof of concept, we focused on chemoreceptors that sense short-chain carboxylic acids in *Pseudomonas* species. These strains are widely distributed^26^ and encode a large repertoire of chemoreceptors^27^. Short-chain carboxylic acids are widespread metabolites in soils^28^ and host-associated environments^29,30^, and are also produced as fermentation products by microbial communities^31^, where they serve as key carbon sources and ecological signals. Using this approach, we successfully identified chemoreceptors responsible for sensing these compounds, providing a simple and scalable framework to map chemoreceptors to metabolic ligands.

### Construction of a cell population with chemoreceptor gene library

To construct a chemoreceptor gene library, we collected 24 *Pseudomonas* strains (**Table S1**). These strains were predominantly isolated from soils and plant-associated environments and encode an average of 41 chemoreceptors per genome. Their chemoreceptors span a wide structural diversity in predicted ligand-binding domains (LBDs) (**Fig.S1**). We extracted their genomic DNA and amplified a total of 975 unique chemoreceptor genes after removing 8 duplicate sequences (**Fig.1A**). These genes were identified based on the annotation “methyl-accepting chemotaxis proteins” in the JGI/IMG database^32^. The complete gene list, including annotation, genomic features, LBD classification, and nucleotide and amino acid sequences, is provided (**Supplementary Table 1**).

**Figure 1:**
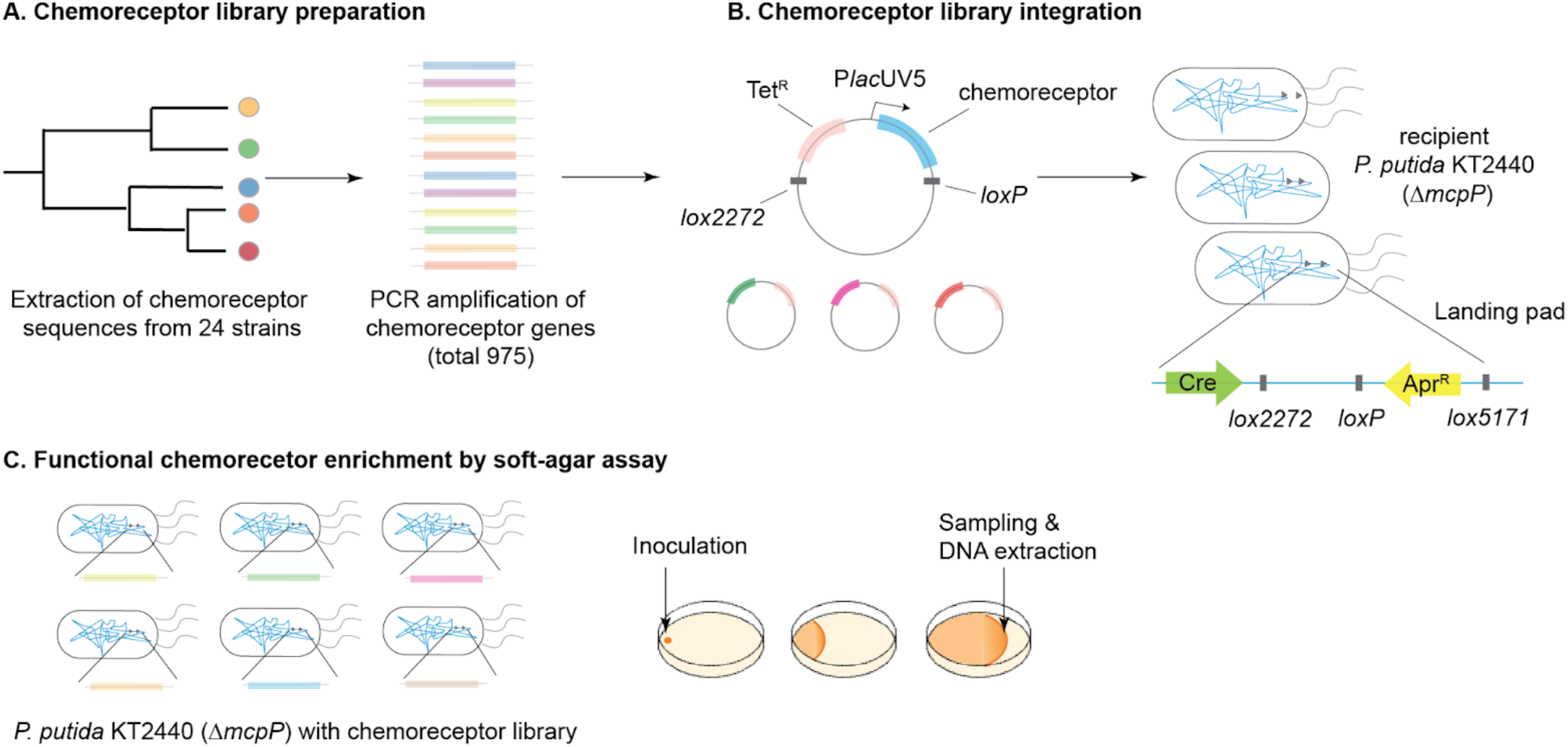
Construction and screening of a pooled chemoreceptor library. (**A**) Chemoreceptor genes were collected from 24 *Pseudomonas* strains and amplified by PCR to generate a library of 975 sequences. (**B**) The pooled chemoreceptor library was cloned under the control of an IPTG-inducible P*lac*UV5 promoter and chromosomally integrated into *Pseudomonas putida* KT2440 *ΔmcpP* (strain MC105) via CRAGE-mediated recombination at a defined landing pad. (**C**) The resulting library population (LS18) was subjected to soft-agar assays for functional enrichment of chemoreceptors based on chemotactic expansion. Cells were collected from the leading edge of expanding populations, followed by DNA extraction and sequencing to quantify enrichment.

The amplified products were pooled at equimolar concentrations and cloned into a shuttle vector (pST003) downstream of an IPTG-inducible promoter (**Fig.1B**). The pooled chemoreceptor library was then chromosomally integrated into a domesticated *Pseudomonas putida* KT2440 *ΔmcpP* (strain MC105) using the CRAGE system^33^ that mediates a targeted recombination, yielding a population expressing a chemoreceptor library (LS18). This population was subsequently used for functional enrichment in soft-agar assays (**Fig.1C**). We selected *P. putida* KT2440 as a heterologous host, because it is a well-characterized plant-root colonizer^34,35^ with flagellar motility and versatile metabolism^36–39^. The *mcpP* deletion, created by Cpf1/Cas12a system^40–42^, eliminates a background chemoreceptor activity for short-chain carboxylic acids, including lactate and propionate^43,44^.

The library was integrated at chromosomal locus 3,784,572 between locus tag PP_3346 and PP_3348 without disrupting coding regions. Soft-agar motility assays verified that the genetic engineering process did not alter the motility phenotypes compared to wild-type strain (**Fig.S2**). Amplicon sequencing from the extracted DNA of a library cell population (LS18) confirmed successful chromosomal integration of 813 chemoreceptor genes out of 975 targets (**Fig.S3, Supplementary Table 1**). Genes that were not recovered were likely lost during library construction steps, including amplification, assembly, or transformation.

### Development of soft-agar enrichment assay

To screen chemoreceptors for interactions with short-chain carboxylic acids, we first tested a soft-agar assay containing 40 mM glycerol and 1 mM lactate as a reference condition. When inoculated at the edge of a soft-agar plate, the wild-type *P. putida* KT2440 exhibited outward expansion via chemotaxis in the presence of lactate, forming a visible ring that propagated at a consistent rate (**Fig.2AB**). This chemotactic expansion was mediated by the chemoreceptor McpP, as the *mcpP* deletion mutant (MC105) showed a reduced expansion rate compared to the wild-type strain (**Fig.2C**: second bar), consistent with previous reports^43,44^. Integration of the chemoreceptor library into *mcpP*-background restored an expansion rate to a level comparable to the wild type (**Fig.2C**: third bar). Induction of library expression with IPTG further increased the expansion rate beyond that of the wild type (**Fig.2C**: fourth bar). Representative images illustrating these differences are shown in **Fig.2D**. Similar results were observed when propionate was used in place of lactate (**Fig.2E**). Because the library strain lacks the native *mcpP* gene, the increased expansion upon induction suggests that functional chemoreceptors within the library mediate chemotactic responses to these compounds.

**Figure 2:**
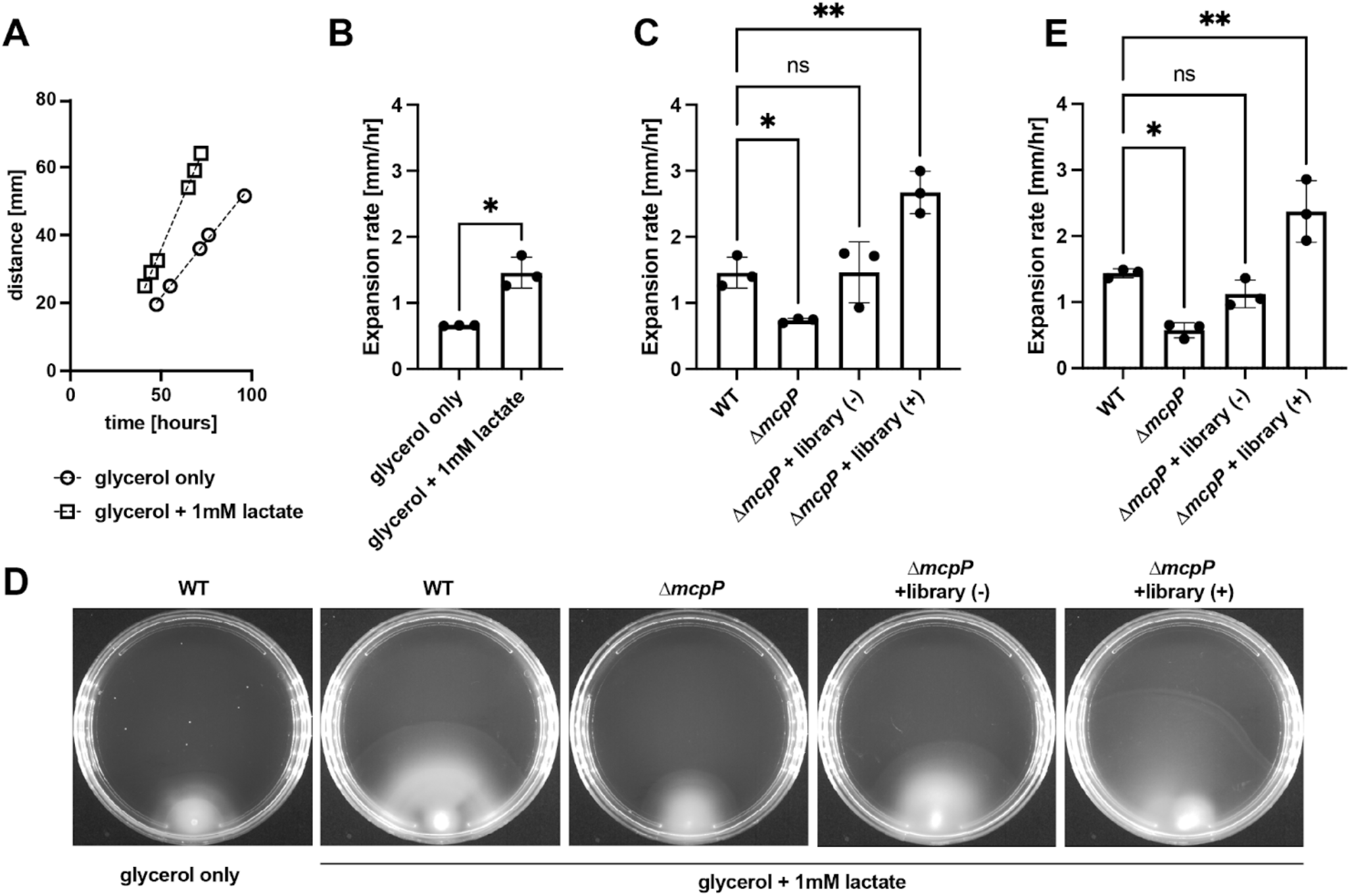
Soft-agar assay to enrich functional chemoreceptors. (**A**) Time-course of colony expansion for *Pseudomonas putida* KT2440 on soft-agar plates containing glycerol alone or glycerol supplemented with 1 mM lactate. (**B**) Quantification of expansion rates under glycerol-only and glycerol + lactate conditions. (**C**) Expansion rates of wild-type (WT), *ΔmcpP* (MC105), and *ΔmcpP* strains carrying the chemoreceptor library (LS18) without (−) or with (+) IPTG induction in the presence of lactate. (**D**) Representative images of colony expansion under the indicated conditions. (**E**) Expansion rates measured in the presence of propionate, showing similar trends as observed with lactate. Data are shown as mean ± s.d. (n = 3 biological replicates). Statistical significance was assessed using a two-tailed Welch’s t-test (B) and ordinary one-way ANOVA (C, E); ns, not significant; *P < 0.05; **P < 0.01.

### Analysis of enriched chemoreceptors

To identify enriched chemoreceptors, we collected cells from the leading edge of expanding populations when they reached near the edge of the plate under induced conditions. In parallel, cells were grown in well-mixed liquid cultures with the same medium composition as the soft-agar assay but without agar and selection for chemotactic behavior (**Fig.3A**). Genomic DNA was extracted from both conditions, and amplicons targeting the library integration site were sequenced using the PacBio platform. Enrichment scores were calculated as the log_2_ ratio of normalized read counts in soft-agar samples relative to those in liquid culture, allowing identification of enriched chemoreceptors.

**Figure 3:**
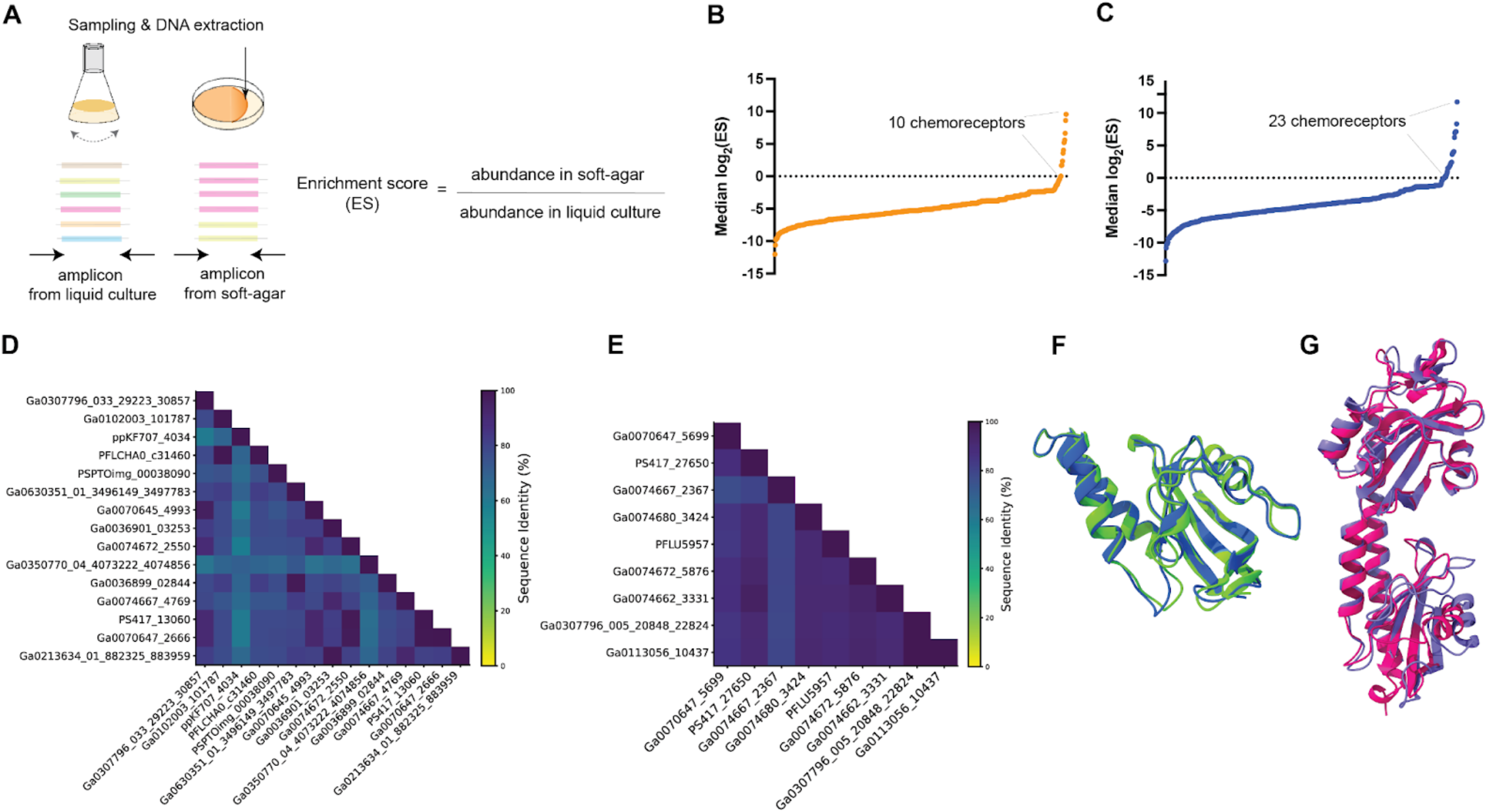
Identification and characterization of enriched chemoreceptors. **(A)** Schematic of enrichment score calculation. Cells from soft-agar and liquid culture conditions were collected, and chemoreceptor abundance was quantified by amplicon sequencing. Enrichment scores (ES) were calculated as the log_2_ ratio of normalized read counts in soft-agar relative to liquid culture. (**B–C**) Ranked median log_2_ enrichment scores of chemoreceptors under lactate (B) and propionate (C) conditions. Chemoreceptors with median log_2_(ES) > 0 were defined as enriched (dashed line), identifying 10 chemoreceptors in lactate and 23 in propionate, with 9 shared between conditions. (**D–E**) Pairwise sequence identity matrices of ligand-binding domains (LBDs) for Group I (D) and Group II (E) chemoreceptors, showing sequence identity ranging from 58.0–99.2 % for Group I and 76.7–99.6 % for Group II. (**F**) Structural alignment of a representative Group I chemoreceptor (PS417_13060) with McpP, showing high structural similarity (RSMD: 0.638 Å). (**G**) Structural alignment of a representative Group II chemoreceptor (PS417_27650) with PacF, showing structural conservation (RSMD: 1.135 Å) despite low sequence identity between them (29.6 %).

The analysis identified 24 enriched chemoreceptors (defined by a median log_2_ enrichment score > 0), including 10 in lactate (**Fig.3B**) and 23 in propionate (**Fig.3C**) with 9 shared in both conditions (**Table 1-2**). Notably, these chemoreceptors are grouped into two groups based on sequence and LBD architecture. Group I chemoreceptors are 544 amino acids in length with minor variation and contain single or double Cache_2 domains (**Table 1**), whereas Group II chemoreceptors are 658 amino acids in length and contain Cache_3–Cache_2 domains (**Table 2)**. Sequence identity analysis of LBDs revealed that Group I chemoreceptors share 58.0 – 99.2 % identity (**Fig.3D**), while Group II chemoreceptors exhibit 76.7 – 99.6 % identity (**Fig.3E**), indicating moderate to high sequence identity within each group.

**Table 1:** Group I chemoreceptors enriched in soft-agar assays and their similarity to McpP.

| Strain origin | Gene ID | Locus tag | Amino acid length | LBD | Enriched in | RSMD to McpP (Å) | Sequence Identity to McpP (%) |
| --- | --- | --- | --- | --- | --- | --- | --- |
| *Pseudomonas marginalis* NCPPB 667 | 2640002922 | Ga0074672_2550 | 544 | dCache_2 | lactate | 0.78 | 77.1 |
| *Pseudomonas furukawai* KF707 | 2532701525 | ppKF707_4034 | 544 | sCache_2 | propionate | 0.719 | 67.9 |
| *Pseudomonas protegens* CHA0 | 2555670089 | PFLCHA0_c31460 | 545 | sCache_2 | propionate | 0.647 | 76.3 |
| *Pseudomonas syringae* pv. tomato DC3000 | 2508869730 | PSPTOimg_00038090 | 541 | dCache_2 | propionate | 0.801 | 75.6 |
| *Pseudomonas oleovorans* NCTC 10692 | 8035396207 | Ga0350770_04_4073222_4074856 | 544 | dCache_2 | propionate | 0.695 | 65.6 |
| *Pseudomonas chlororaphis aureofaciens* 30-84 | 2597858032 | Ga0036899_02844 | 544 | dCache_2 | propionate | 0.675 | 76.3 |
| *Pseudomonas kilonensis* DSM 13647 | 2640374745 | Ga0074667_4769 | 544 | dCache_2 | propionate | 0.578 | 72.5 |
| *Pseudomonas simiae* WCS417 | 2586031246 | PS417_13060 | 544 | dCache_2 | propionate | 0.638 | 78.6 |
| *Pseudomonas lurida* LMG 21995 | 2616887541 | Ga0070647_2666 | 544 | dCache_2 | propionate | 0.713 | 79.4 |
| *Pseudomonas fluorescens* FW300-N2E2 | 2875505803 | Ga0213634_01_882325_883959 | 544 | dCache_2 | propionate | 0.678 | 74.8 |
| *Pseudomonas fluorescens* 2-79 | 8011198331 | Ga0307796_033_29223_30857 | 544 | sCache_2 | lactate & propionate | 0.75 | 79.4 |
| *Pseudomonas* sp. 397MF | 2643584364 | Ga0102003_101787 | 544 | sCache_2 | lactate & propionate | 0.664 | 76.3 |
| *Pseudomonas chlororaphis piscium* ATCC 17411 | 8088465072 | Ga0630351_01_3496149_3497783 | 544 | dCache_2 | lactate & propionate | 0.67 | 77.1 |
| *Pseudomonas fluorescens* ATCC 13525 | 2617959847 | Ga0070645_4993 | 544 | dCache_2 | lactate & propionate | 0.695 | 78.6 |
| *Pseudomonas fluorescens* Q8r1-96 | 2597876007 | Ga0036901_03253 | 544 | dCache_2 | lactate & propionate | 0.942 | 75.6 |

**Table 2:** Group II chemoreceptors enriched in soft-agar assays and their similarity to PacF.

| Strain origin | Gene ID | Locus tag | Amino acid length | LBD | Enriched in | RSMD to PacF (Å) | Sequence Identity to PacF (%) |
| --- | --- | --- | --- | --- | --- | --- | --- |
| *Pseudomonas fluorescens* SBW25 | 649639360 | PFLU5957 | 658 | Cache_3-Cache_2 | propionate | 1.199 | 29.9 |
| *Pseudomonas marginalis* NCPPB 667 | 2640006248 | Ga0074672_5876 | 658 | Cache_3-Cache_2 | propionate | 1.259 | 29.1 |
| *Pseudomonas extremorientalis* LMG 19695 | 2640452441 | Ga0074662_3331 | 658 | Cache_3-Cache_2 | propionate | 1.096 | 29.7 |
| *Pseudomonas fluorescens* 2-79 | 8011195395 | Ga0307796_005_20848_22824 | 658 | Cache_3-Cache_2 | propionate | 1.143 | 29.3 |
| *Pseudomonas synxantha* DSM 18928 | 2662690122 | Ga0113056_10437 | 658 | Cache_3-Cache_2 | propionate | 1.13 | 29.4 |
| *Pseudomonas lurida* LMG 21995 | 2616890574 | Ga0070647_5699 | 658 | Cache_3-Cache_2 | lactate & propionate | 1.095 | 28.3 |
| *Pseudomonas simiae* WCS417 | 2586034105 | PS417_27650 | 658 | Cache_3-Cache_2 | lactate & propionate | 1.135 | 29.6 |
| *Pseudomonas kilonensis* DSM 13647 | 2640372343 | Ga0074667_2367 | 658 | Cache_3-Cache_2 | lactate & propionate | 1.086 | 28.9 |
| *Pseudomonas poae* LMG 21465 | 2640299616 | Ga0074680_3424 | 658 | Cache_3-Cache_2 | lactate & propionate | 1.207 | 29.4 |

Sequence analysis revealed that the LBDs of Group I share 65.6 – 79.4 % sequence identity with McpP (**Table 1**), which was deleted from *P. putida* KT2440 before library integration. Although sequence identity is moderate, all Group I chemoreceptors conserve key residues required for ligand interaction^23,45^ (**Fig.S4**). Structural analysis of LBDs further showed close alignment with McpP, with root mean square deviation (RMSD) values ranging from 0.58 to 0.80 Å (**Table 1**). A representative structural alignment between McpP and a chemoreceptor from *Pseudomonas simiae* WCS417 (PS417_13060) is shown (**Fig.3F**). Together, these results indicate that Group I chemoreceptors are conserved relatives of McpP.

In contrast, Group II chemoreceptors possess a Cache_3–Cache_2 domain architecture, a variant of the dCache domain^46^, which remains poorly characterized within the Cache family. A recent study^47^ identified PacF, a chemoreceptor from *Pectobacterium atrosepticum*, as a representative of this domain class. Notably, despite low sequence identity of LBDs to PacF (28.3 – 29.9 %) (**Table 2**), Group II chemoreceptors retain conserved ligand-interacting residues with PacF (**Fig.S5**). Structural comparisons of LBDs revealed close alignment of Group II chemoreceptors against PacF, with RMSD values ranging from 1.09 to 1.26 Å (**Table 2**). A representative alignment between PacF and a chemoreceptor from *Pseudomonas simiae* WCS417 (PS417_27650) is shown (**Fig.3G**). These results suggest that Group II chemoreceptors represent a sequentially diverged yet structurally conserved family of the Cache_3–Cache_2 class.

A key distinction from previous work^47^ is that the Cache_3–Cache_2 chemoreceptors identified in this study were enriched using C3 ligands, including lactate and propionate, whereas PacF was reported to respond specifically to the C1 molecule formate. To assess their function, we isolated representative chemoreceptors from our screen and tested their chemotactic responses individually. All four isolates exhibited enhanced chemotactic expansion in the presence of C3 carboxylic acids (**Fig.4A–C**), with clear ring formation (**Fig.4E**), including lactate, propionate, and pyruvate. In contrast, no enhanced expansion was observed in response to formate (**Fig.4D**). However, ring formation was observed in a subset of isolates (PFLU5957 and Ga0074667_2367) (**Fig.4F**), indicating weak and variable chemotactic responses. Growth measurements showed that *P. putida* KT2440 utilized formate at levels comparable to other carbon sources (**Fig.S6**), suggesting that the lack of enhanced expansion is not due to limited metabolism. Together, these results demonstrate that Cache_3–Cache_2 chemoreceptors in *Pseudomonas* primarily mediate chemotaxis toward short-chain C3 carboxylic acids, thereby expanding the functional scope of this domain.

**Figure 4:**
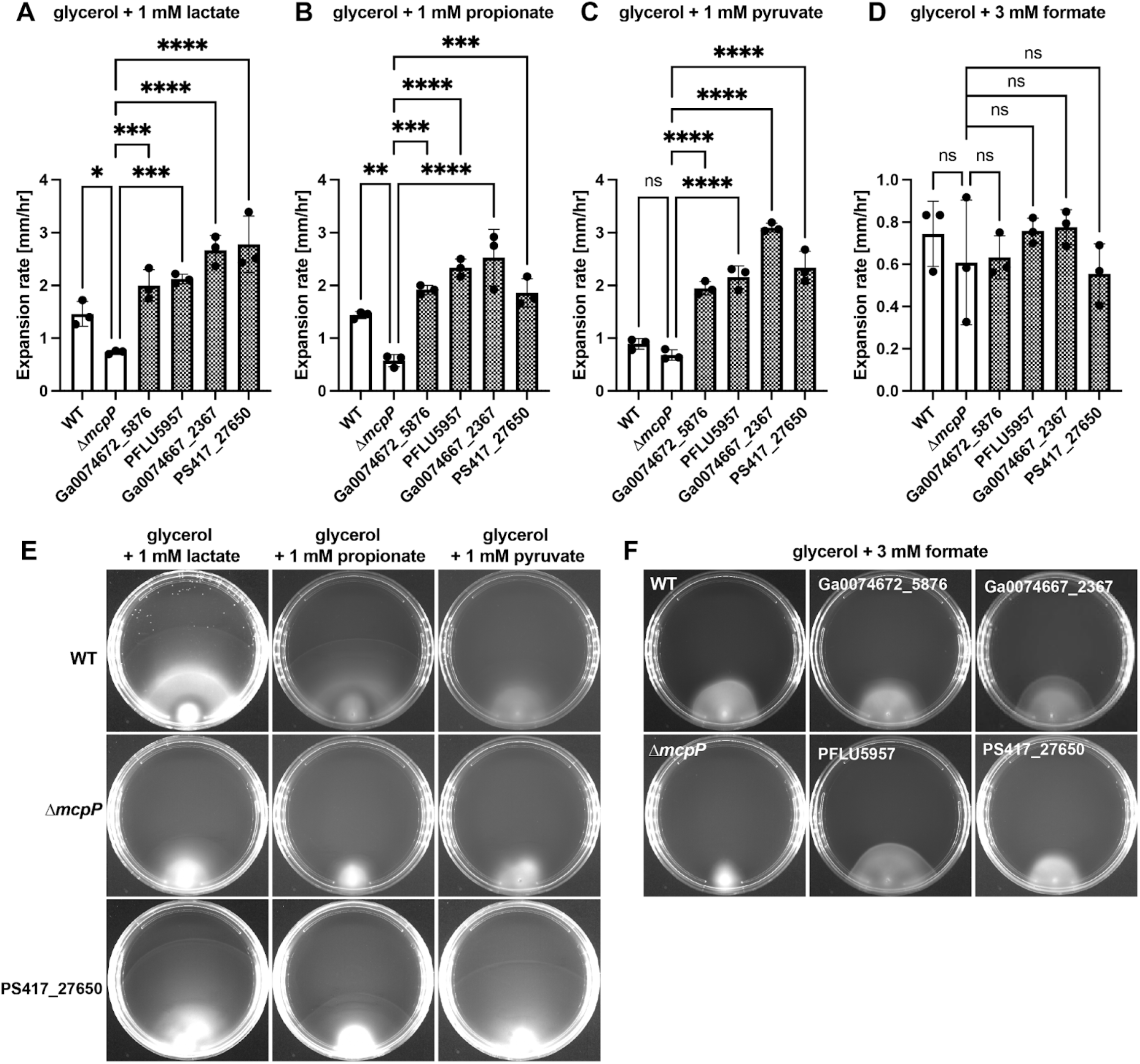
Functional characterization of selected chemoreceptors with Cache_3-Cache_2 domains. (**A–D**) Expansion rates in soft-agar assays containing 40 mM glycerol supplemented with 1 mM lactate (A), 1 mM propionate (B), 1 mM pyruvate (C), or 3 mM formate (D). Wild type (WT), *ΔmcpP* (strain MC105), and 4 complemented strains expressing individual Cache_3-Cache_2 domain chemoreceptors from different strain origins were tested (locus tags are indicated; see **Table 2**). Complemented strains were tested with 1 mM IPTG. (**E**) Representative images of soft-agar plates for WT, Δ*mcpP*, and a strain complemented with PS417_27650 across lactate, propionate, and pyruvate conditions. (**F**) Representative images of soft-agar plates for WT, Δ*mcpP*, and complemented strains in formate conditions. Data are shown as mean ± s.d. (n = 3 biological replicates). Statistical significance was assessed using ordinary one-way ANOVA with multiple comparisons; ns, not significant; *P < 0.05; **P < 0.01; ***P < 0.001; ****P < 0.0001.

Our structural and molecular dynamics analyses indicate that differences in ligand-binding pocket architecture may underlie this shift in ligand specificity. Specifically, the predicted ligand access tunnel of the representative chemoreceptor (PS417_27650) exhibits larger volume than that of PacF at the membrane-distal module of the dCache domain (**Fig.5AB**). In addition, analysis of the ligand-binding pocket shaped by key residues revealed a ∼2.6-fold increase in volume compared to PacF (**Fig.5CD**), consistent with accommodation of larger C3 carboxylic acids. While key pocket residues are conserved between PS417_27650 and PacF, substitution of a smaller valine (V130) in PS417_27650 for a bulkier isoleucine (I132) in PacF likely contributes to the enlarged pocket by permitting a more open local structure. Molecular dynamics simulations further suggest that sequence variations alter the flexibility of a loop region at the membrane-distal module of the dCache domain, resulting in a more stabilized pocket in the PS417_27650 compared to PacF (**Fig.S7AB**). These structural features are associated with longer ligand residence time for lactate relative to PacF (**Fig.S7C**). Together, these results support a model in which expanded access tunnels and enlarged binding pockets, coupled with altered flexibility, facilitate recognition of larger C3 ligands.

**Figure 5:**
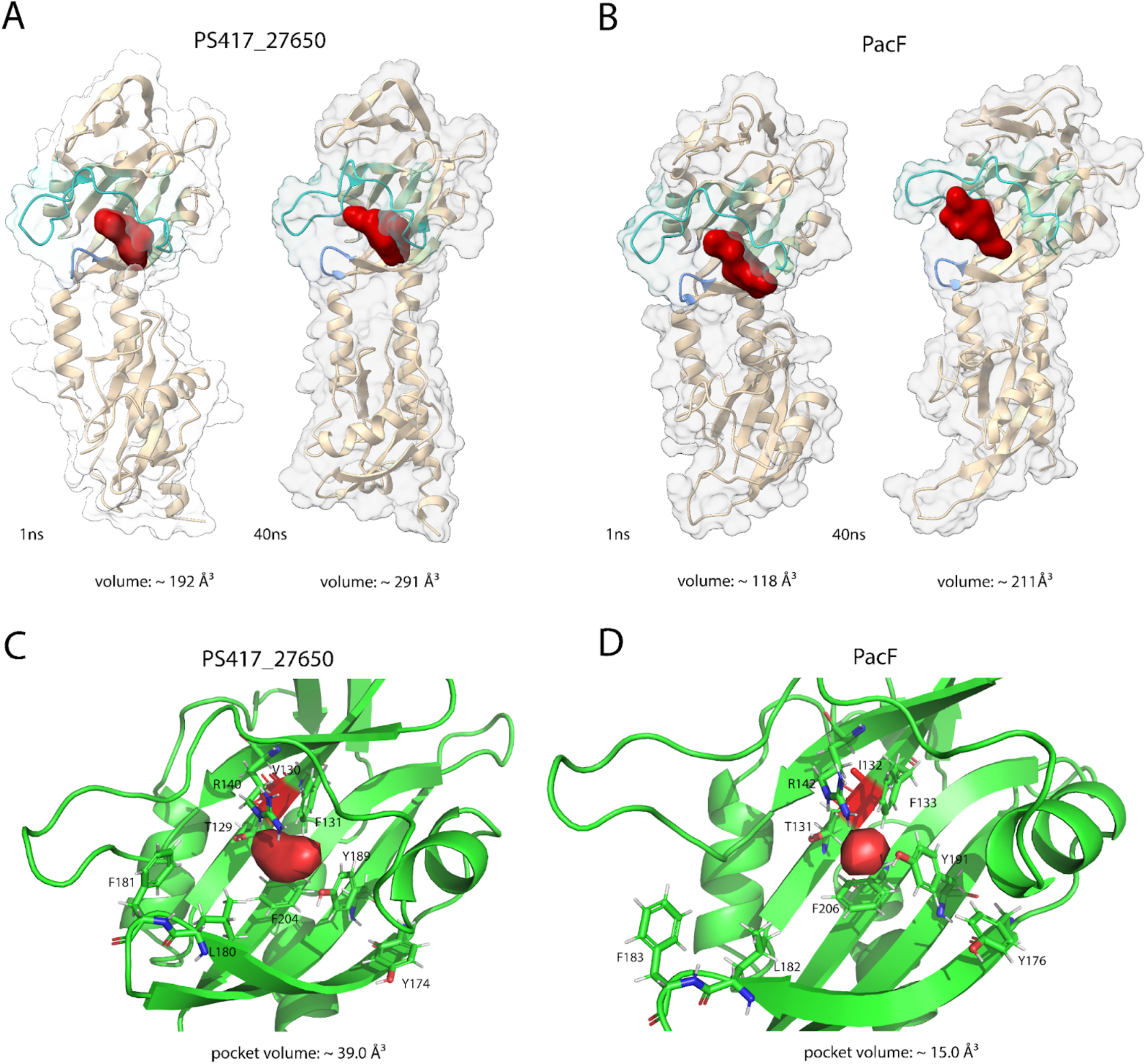
Structural comparison of ligand access tunnels and binding pockets. (**A–B**) Representative structural snapshots of PS417_27650 (A) and PacF (B) at 1 ns and 40 ns from molecular dynamics simulations. The ligand access tunnel is shown in red, and estimated tunnel volumes are indicated below. (**C–D**) Close-up views of the ligand-binding pockets at 40 ns in PS417_27650 (C) and PacF (D), highlighting the available pocket space (red) defined by conserved residues R140/142, F131/133, Y189/191, F204/206, and T129/131, corresponding to key ligand-interacting residues previously reported for PacF^47^. PS417_27650 exhibits an approximately 2.6-fold increase in pocket volume compared to PacF. These results are consistent with accommodation of larger C3 carboxylic acids in PS417_27650 relative to the C1 formate sensed by PacF.

## Discussion

In this study, we developed a pooled screening approach to identify functional chemoreceptors in *Pseudomonas* strains that sense short-chain C3 carboxylic acids. This approach addresses key limitations of existing methods, including the difficulty of genetic mutant construction in non-model bacteria and challenges in protein folding and purification associated with in vitro assays and chimeric receptor expression. Heterologous expression of chemoreceptors in a closely related host strain likely facilitates proper folding, membrane localization, array formation and coupling to downstream signaling pathways. Consistent with this, our screening results revealed enrichment of functional chemoreceptors originating from diverse strains (**Table 1-2**).

Using this platform, we discovered a group of chemoreceptors with Cache_3-Cache_2 domains in *Pseudomonas* strains that sense C3 carboxylic acids, including lactate, propionate, and pyruvate. These chemoreceptors share low sequence identity (∼29 %) in their LBDs with the recently characterized PacF, which senses the C1 molecule formate, yet they retain conserved structural features and key motifs. Our computational analyses suggest that amino acid substitutions at key positions contribute to an enlarged ligand-binding pocket, thereby enabling recognition of larger C3 ligands in the identified chemoreceptors. This is consistent with the rapid evolvability of chemoreceptors to recognize novel ligand combinations^23^, facilitating bacterial adaptation to new environmental niches.

Several aspects of the current approach can be further optimized to expand its applicability. First, the host strain *Pseudomonas putida* KT2440 retains 27 endogenous chemoreceptors, and only a single chemoreceptor (*mcpP*) was deleted in this study. As residual background activity from native chemoreceptors can interfere with heterologous chemoreceptor functions in the library, future work will require deletion of multiple or all endogenous chemoreceptor to improve screening specificity. Second, while the use of soft-agar assays provides a simple and scalable setup, they require ligand consumption and limits the detection of compounds that are poorly metabolized. Incorporating alternative assays, such as high-throughput capillary assays^48,49^, could enable broader ligand screening. Finally, a subset of chemoreceptors was unselected after the screening assay despite being successfully cloned into the library and sharing similar sequence identities with enriched ones (**Table S5**), suggesting biases introduced during experimental steps. This limitation could be mitigated by improving library uniformity, optimizing sampling protocols and sequencing library preparation.

Although further refinement is required, this approach provides a scalable framework for high-throughput functional characterization of chemoreceptors. As *Pseudomonas* species encode large and diverse chemoreceptor repertoires, the findings will provide a foundation for annotating chemoreceptors and other sensor proteins across diverse bacterial lineages. Moreover, the generated data offer valuable training datasets for computational models, which remain limited in their ability to accurately predict ligand specificity from structural information alone^50,51^. A comprehensive mapping of ligand-chemoreceptor interactions will establish sequence–structure–function relationships in sensor proteins and has the potential to accelerate the development of novel biosensors^52,53^.

### Bacterial strains

We used a wild-type *Pseudomonas putida* KT2440 (ATCC 47054) as a host of a chemoreceptor gene library. The gene library was prepared from the following strains as listed in **Table S1**: *Pseudomonas fluorescens* (ATCC 1352), *Pseudomonas protegens* CHA0 (DSM 19095), *Pseudomonas corrugata* (DSM 7228), *Pseudomonas saponiphila* (DSM 9751), *Pseudomonas marginalis* NCPPB 667 (DSM 13124), *Pseudomonas thivervalensis* (DSM 13194), *Pseudomonas poae* LMG 21465 (DSM 14936), *Pseudomonas koreensis* (DSM 16610), *Pseudomonas oleovorans* NCTC 10692 (DSM 1045), *Pseudomonas taiwanensis* (DSM 21245), *Pseudomonas synxantha* (DSM 18928), *Pseudomonas furukawaii* KF707 (DSM 10086), *Pseudomonas kilonensis* (DSM 13647), *Pseudomonas flourescens* Q8-R1, *Pseudomonas fluorescens* 2-79, *Pseudomonas fluorescens* SBW25, *Pseudomonas chlororaphis piscium* (ATCC 17411), *Pseudomonas chlororaphis aureofaciens* 30-84, *Pseudomonas simiae* WCS417, and *Pseudomonas* sp. 397MF.

### Landing pad integration

The wild-type *Pseudomonas putida* KT2440 was domesticated by integrating a landing pad compatible with CRAGE^33,54^. The resulting strain was designated MC34 (**Table S2**). First, the landing pad region, including Cre recombinase and the recombination sites *lox2272, loxP*, and *lox5171*, was amplified from plasmid pEC387 using primer sets with extensions recognized by Tn5. Then, the amplified fragment was chromosomally integrated using the EZ-Tn5 Transposase Kit (LGC Biosearch Technologies; TNP92110) and transformants were selected on LB plates supplemented with kanamycin. Approximately 10 colonies were screened to determine integration sites by PacBio sequencing. A clone with insertion at chromosomal position 3,784,572, located at the intergenic region between locus tags PP_3346 and PP_3348 (74 bp downstream of PP_3346 end codon and 321 bp upstream of PP_3348 start codon), was selected for further experiments.

### Construction of *mcpP* deletion mutant

Deletion of the *mcpP* (locus_tag: PP_2861) was performed by using the Cpf1/Cas12a system^40,41^. We used two plasmids (**Table S3**): pORTMAGE-Pa1 (Addgene #138475), which carries *recT, mutL* and *GenR*, as a recombineering plasmid, and pTE433-beta, which carries *Cpf1/Cas12a* and *KanR* as a cpf1-gRNA plasmid. The latter one was modified in this study as follows. First, we replaced the kanamycin resistance gene with an apramycin resistance gene, because our domesticated *P. putida* strain (MC34) already carried kanamycin resistance. Second, a guide RNA targeting *mcpP* was inserted, resulting in the plasmid pTE433-mcpP. The guide RNA sequence was designed by identifying a 21 bp target site adjacent to a 5′-TTTN-3′ PAM sequence within the *mcpP* coding region: 5’-TCCGGCGGCCTGTTCAGCGGT-3’. The plasmid assemblies were performed using the Gibson Assembly HiFi kit (New England Biolabs: E2621), and *Escherichia coli* TOP10 (Thermo Fisher Scientific: c4040).

For recombineering, the plasmid pORTMAGE-Pa1 was first transformed into MC34 by electroporation. The transformant was then inoculated into LB medium supplemented with 50 µg/ml gentamicin and incubated overnight at 30 °C with the shaking at 200 r.p.m. Induction of recombinase expression was carried out by adding 10 mM 3-methyl-benzoate (Sigma: T36609), followed by shaking for an additional 30 minutes. Cells were then washed 3 times with ice-cold 10 % glycerol, resuspended in 500 µl 10% glycerol, and 50 µL of the cells was electroporated with 5 µl of 100 µM donor single-strand DNA (200 bases, Integrated DNA Technologies, **Table S4**) and 200 ng of pTE433-mcpP. The cells were recovered at 30 °C for 3–4 hours in the shaking incubation at 120 r.p.m. and subsequently selected on plates containing 200 µg/ml apramycin. Colony PCR was performed on the resulting colonies to confirm gene deletion with mcpP_fwd and mcpP_rev primers (**Table S4**). We selected one of the obtained mutants (MC63) for subsequent experiments. We confirmed that both the recombineering plasmid and CRISPR plasmid were cured from the mutant by checking the sensitivities to gentamicin and apramycin. Prior to the library construction, we inserted *mVenus* fluorescein gene into MC63 for possible utility of fluorescent assays. For this, we used a plasmid pP5171-mVenus which carries *mVenus* flanked by *lox5171* and *loxP* with apramycin resistance gene. The plasmid was introduced into MC63 by conjugation with *E. coli* WM3064, resulting in a strain (MC105) that was subsequently used for library integration.

### Chemoreceptor library construction and integration

Genomic DNAs were extracted from 24 *Pseudomonas* strains using Quick-DNATM Fungal/Bacterial Miniprep Kit (Zymo Research: D6005). Chemoreceptor coding sequences were amplified by plate-based PCR from genomic DNA using the primers listed in **Supplementary Table 2** using Platinum™ SuperFi II (Thermo Fisher Scientific: 12368). Primers were designed to anneal 25 bp upstream and downstream of the start and stop codons, with additional 25 bp extensions for cloning. All amplicons were purified (Promega: A9281, Cytiva: 7700) and quantified using the Qubit BR assay kit (Invitrogen: Q32853). Equimolar amounts of individual amplicons were pooled together. A total of 80 ng of the pooled amplicon DNA was cloned into 70 ng of pST003 vector digested with XhoI (New England Biolabs, R0146) and XbaI (New England Biolabs, R0145), using NEBuilder HiFi DNA Assembly Master Mix (New England Biolabs: E2621). The pST003 vector enables expression of inserted chemoreceptor genes under the control of the P*lac*UV5 promoter with IPTG induction. This plasmid was derived from pW38^54^ with the following modifications: (i) replacement of the kanamycin resistance gene with a tetracycline resistance gene; (ii) deletion of the T7 RNA polymerase gene; and (iii) insertion of a Shine–Dalgarno sequence, multiple cloning site, and T7 terminator downstream of lacO.

To assess the cloning efficiency, a negative control was prepared in parallel without the addition of the pooled amplicons. Both DNA samples were bead purified separately using two volumes of Mag-Bind® TotalPure NGS (Omega Bio-tek Inc: M1378), and the purified DNA was electroporated into TransforMax™ EC100D™ *pir*^*+*^ Electrocompetent *E. coli* (Biosearch Technologies: ECP09500). To titer the transformation efficiency of the pooled library, ten-fold serial dilutions of the transformed cells were inoculated on LB agar plate containing 10 μg/ml tetracycline. An estimated 8 × 10^5^ cells (∼800-fold the library size) were inoculated into 150 ml LB containing 10 μg/ml tetracycline and grown overnight at 30 °C and 200 r.p.m. Cells were harvested and the plasmid library was purified using ZymoPURE II Plasmid Midiprep Kit (Zymo Research: D4209). The plasmid library was transformed into a strain (MC105) by electroporation and the cassette exchange was mediated by CRAGE recombination^54^. The cells were then spread to a 24.5 cm square petri dish (Corning: 07-200-600), and all colonies were collected into 20 ml LB medium containing 10% glycerol, and stored at -80 °C. This served as the chemoreceptor library (LS18) that we used for screening experiments. Of the 975 chemoreceptor genes targeted for amplification, 813 were successfully introduced into MC105. The genes that were not successfully introduced were likely lost during gene amplification, assembly, or transformation. Complemented strains with Cache_3-Cache_2 chemoreceptors were generated by reintroducing individual cloned chemoreceptors into MC105.

### Growth rate measurement

Each experiment consisted of three steps: seed culture, pre-culture, and experimental culture. For the seed culture, a single colony from a freshly streaked LB agar plate was inoculated into 3 ml liquid LB medium in a 14 ml plastic tube and incubated overnight at 28 °C in a shaking incubator at 200 r.p.m. Cells from the seed culture were diluted into 2 ml of fresh minimal media containing 1x M9 minimal salts (Gibco: A1374401), 2 mM magnesium sulfate (Sigma: M3409), 0.1 mM calcium chloride (Sigma: 223506), and 10 µM ferrous sulfate (Sigma: F8633), and 40 mM glycerol (Sigma: G5516). The overnight pre-culture was washed twice using 1x M9 minimal salts, then subsequently diluted to an initial OD_600_ of 0.005 in the fresh minimal M9 medium. For the experimental cultures, the medium was supplemented with either 5 mM glycerol or 5 mM glycerol combined with 5 mM of lactate (Sigma: 71720), propionate (Sigma: P1880), pyruvate (Sigma: P5280), or formate (Sigma: 71539) as carbon sources. These cultures were then incubated at 28 °C in the Infinite F200 Pro plate reader (Tecan), where OD_600_ was monitored every 30 min. All experiments were performed with 3 biological replicates.

### Soft-agar assay

Expansion speeds were measured using soft-agar plates containing 1x M9 minimal salts, 2 mM magnesium sulfate, 0.1 mM calcium chloride, and 10 µM ferrous sulfate, and 0.25 % agar (BD Difco: 214040), supplemented with defined carbon sources. The primary condition consisted of 40 mM glycerol supplemented with 1 mM lactate, 1 mM propionate, 1 mM pyruvate, or 3 mM formate. Additional conditions, including 10 mM glucose, 10 mM citrate, and 2.5% LB, were used to assess the effects of genetic engineering (**Fig.S2**). Where indicated, 1 mM IPTG was added to induce expression of the chemoreceptor library or complemented Cache_3-Cache_2 chemoreceptors. For plate preparation, 15 ml of freshly prepared, still-warm soft-agar medium was poured into a 10-cm-diameter Petri dish (Corning: 25373), forming a ∼2-mm-thick agar layer. Plates were allowed to solidify at room temperature for at least 10 min and were freshly prepared prior to each assay. Cell cultures were prepared by following seed culture and pre-culture, except that 0.05% PVP40 (Sigma: PVP40) was added during these steps to prevent cells and flagella from sticking to material surfaces^55^. For the assay, 2 µl of pre-culture diluted to an OD_600_ of 0.5 was spotted onto the edge of each soft-agar plate, and incubated at 28 °C for 2-4 days. Once the expanding population reached a radius of at least 2 cm, expansion was assessed by manual observation. The distance from the inoculation point to the edge of the expanding ring was measured every 3–4 hours over at least three time points. Expansion speeds were calculated by linear regression of colony radius over time. All experiments were performed with three biological replicates.

### Screening assay

A pre-culture of the chemoreceptor library population (LS18) was prepared in M9 medium supplemented with 40 mM glycerol, 1 mM IPTG, and 10 µg/mL tetracycline. For soft-agar assays, 2 µl of pre-culture diluted to an OD_600_ of 0.5 was inoculated onto the edge of soft-agar plates containing 1 mM IPTG and the selected carbon sources: 40 mM glycerol supplemented with 1 mM lactate or 1 mM propionate. Plates were incubated at 28 °C for 2–4 days. Once the expanding colony reached a radius of approximately 5 cm from the inoculation point, cells were collected from the outer expansion ring. In parallel, liquid cultures with the same composition (excluding agar) were prepared by inoculating 2 µL of the pre-culture into liquid medium. Cultures were incubated for 2 days under the same conditions, and cells were harvested. Genomic DNA was extracted from both soft-agar and liquid samples using the Quick-DNA™ Fungal/Bacterial Miniprep Kit (Zymo Research; D6005).

### Sequencing library preparation

The chemoreceptor genes were amplified by a two-step PCR protocol from the extracted DNA. The first PCR reaction was carried out in a total volume of 25 µl for each sample, containing 12.5 µL of KAPA polymerase (Roche: KK2602), 2.5 µL each of primer (M13-tailed_fwd/rev primers, **Table S4**), 10 ng of extracted DNA, and nuclease-free water to a final volume of 25 µl. The PCR conditions were as follows: (1) initial heating at 95 °C for 3 minutes, (2) 20 cycles of 20 seconds at 98 °C, 15 seconds at 65 °C, and 2.5 minutes at 72 °C, (3) a final extension at 72 °C for 5 minutes. After purification by using DNA Clean & Concentrator-5 (Zymo Research: D4014), the resulting amplicons were used for a second PCR with barcoding primers. The second PCR was carried out in a total volume of 25 µl for each sample, containing 12.5 µL of KAPA polymerase (Roche: KK2602), 2.5 µL each of primer (Barcoded M13_bc primers, **Table S4**), 10 ng of first PCR product, and nuclease-free water to a final volume of 25 µl. The PCR conditions were as follows: (1) initial heating at 95 °C for 3 minutes, (2) 2 cycles of 20 seconds at 98 °C, 15 seconds at 60 °C, and 2.5 minutes at 72 °C, (3) 5 cycles of 20 seconds at 98 °C, 15 seconds at 65 °C, and 2.5 minutes at 72 °C, (4) a final extension at 72 °C for 5 minutes. Library preparation followed a PacBio protocol for multiplexed amplicon sequencing using barcoded M13 primers. All PCR products were purified with DNA Clean & Concentrator-5 (Zymo Research: D4014). Final amplicon concentrations were determined using the Qubit High Sensitivity (HS) Assay Kit (Invitrogen: Q32854), and fragment sizes were checked using the High Sensitivity DNA Kit on the Bioanalyzer system (Agilent Technologies: 5067-4626). Sequencing was performed on a PacBio Revio 25M SMRT Cell at QB3 Genomics, UC Berkeley.

### Data analysis

Sequencing reads were demultiplexed based on barcode sequences and mapped to the reference chemoreceptor gene library using a custom library quality control pipeline. Counts per million (CPM) were calculated for each gene in each sample. Enrichment scores were defined as the log_2_ ratio of (CPM + 1) in soft-agar samples to (CPM + 1) in liquid culture samples for each gene and each replicate. Genes with a median CPM of 0 across replicates in liquid culture were excluded.

### Molecular dynamics simulation

All molecular dynamic (MD) simulations were performed using GROMACS 2025.2^56^. Briefly, we selected PS417_27650 as the representative for our isolated Cache_3-Cache_2 chemoreceptors and compared its dynamic behaviors to that of pacF in water and in complex with multiple lactate compounds. The structure of PS417_27650 and pacF ligand binding domains were prepared by Alphafold 3^57^ and subjected to 1 µs energy minimization using OpenMM^58^ to resolve any potential clashes. Lactate parameters were generated using CGenFF server (https://cgenff.com/). In the case of protein-lactate complexes, one lactate molecule is placed at the center of LBD of pacF and PS417_27650, and nine other lactate molecules are placed randomly near the protein. Protein topologies were prepared with the CHARMM36^59^ force field and TIP3P water model. Protein-lactate complexes were solvated using the SPC216 water model and neutralized with Na^+^ and Cl^-^ ions at physiological salt concentration. Energy minimization was performed by steepest descent minimization algorithm with a maximum allowed iteration of 50,000 steps, and convergence threshold < 500 kJ/mol/nm followed by NVT and NPT equilibration. NVT was performed at 300 K using the V-rescale thermostat algorithm^60^ with a coupling constant of 0.1 ps for 100 ps, while NPT was conducted at 300 K using Berendsen thermostat^61^ for 100 ps. Both NVT and NPT utilized a time step of 2 fs, with long-range electrostatics (cutoff distance 1.2 nm) calculated by Particle Mesh Ewald (PME) algorithm^62^ and van der Waals interactions calculated by cut-off method (1.2 nm). MD simulations were run for 50 ns using the C-rescale^63^ for pressure coupling and V-rescale for temperature coupling, maintaining a temperature of 300 K and a 2 fs time step. Each protein-ligand complex system was simulated for 50 ns in at least two replicates, ensuring a more robust analysis of protein-ligand interactions. PS417_27650 and pacF were also simulated in water for 50 ns to make sure the observed behaviors of the proteins are consistent. Protein–ligand interactions were analyzed from the resulting trajectories. The distance of the center of mass (COM) of the lactate in the pocket to the COM of the key residue arginine was used to estimate the dissociation times of lactate from the complex.

## Acknowledgement

We thank Dr. Ian Blaby, Dr. Thomas Eng and Dr. Rekha Seshadri for helpful discussions and technical suggestions. We also thank the Joint Genome Institute and QB3 Genomics Center at UC Berkeley for library construction and sequencing. This work was supported by the Laboratory Directed Research and Development (LDRD) Program at Lawrence Berkeley National Laboratory under U.S. Department of Energy project identifier No. LB24023 (H.O., S.Y., T.H.). Work conducted by the U.S. Department of Energy Joint Genome Institute, a DOE Office of Science User Facility, is supported by the Office of Science of the U.S. Department of Energy under Contract No. DE-AC02-05CH11231 (H.O., D.M., N.G., Z.Y.W., D.R., Y.Y., N.M., T.H.). Additional support was provided by the Bioimaging Program (S.Y., Y.Y., T.H.) and the Biosystems Design Program (N.P.) of the U.S. Department of Energy, Office of Biological and Environmental Research, under Contract No. DE-AC02-05CH11231 and DE-SC0023091.

## Author contribution

T.H. and H.O. designed the study. H.O., S.Y., and T.H. performed the experiments, with contributions from N.G., who assisted in the construction of the chemoreceptor gene library. H.O. and T.H. analyzed the data. D.M. performed structural analyses. S.Y., Z.Y.W., N.P., and D.R. contributed to plasmid construction. T.H., Y.Y., and N.M. acquired funding. T.H. supervised the study. T.H. wrote the manuscript with contributions from H.O. and D.M. All authors reviewed and approved the final version of the manuscript.

## References

1. Matilla, M. A. & Krell, T. The effect of bacterial chemotaxis on host infection and pathogenicity. FEMS Microbiol. Rev. 42, fux052 (2018).

2. Liu, Y. et al. Root colonization by beneficial rhizobacteria. FEMS Microbiol. Rev. 48, fuad066 (2024).

3. Zhou, B., Szymanski, C. M. & Baylink, A. Bacterial chemotaxis in human diseases. Trends Microbiol. 31, 453–467 (2023).

4. Gumerov, V. M., Andrianova, E. P. & Zhulin, I. B. Diversity of bacterial chemosensory systems. Curr. Opin. Microbiol. 61, 42–50 (2021).

5. Matilla, M. A., Velando, F., Martín-Mora, D., Monteagudo-Cascales, E. & Krell, T. A catalogue of signal molecules that interact with sensor kinases, chemoreceptors and transcriptional regulators. FEMS Microbiol. Rev. 46, fuab043 (2022).

6. Brunet, M. et al. An atlas of metabolites driving chemotaxis in prokaryotes. Nat. Commun. 16, 1242 (2025).

7. Keegstra, J. M., Carrara, F. & Stocker, R. The ecological roles of bacterial chemotaxis. Nat. Rev. Microbiol. 20, 491–504 (2022).

8. Cole, B. J. et al. Genome-wide identification of bacterial plant colonization genes. PLOS Biol. 15, e2002860 (2017).

9. Aroney, S. T. N., Poole, P. S. & Sánchez-Cañizares, C. Rhizobial Chemotaxis and Motility Systems at Work in the Soil. Front. Plant Sci. 12, 725338 (2021).

10. Wu, X. et al. Metagenomic insights into genetic factors driving bacterial niche differentiation between bulk and rhizosphere soils. Sci. Total Environ. 891, 164221 (2023).

11. Ramoneda, J. et al. Ecological relevance of flagellar motility in soil bacterial communities. ISME J. wrae067 (2024) doi:10.1093/ismejo/wrae067.

12. Yang, C.-X. et al. Plant Exudates Driven Microbiome Recruitment and Assembly Facilitates Plant Health Management. FEMS Microbiol. Rev. fuaf008 (2025) doi:10.1093/femsre/fuaf008.

13. Honda, T. et al. CRAGE-RB-PI-seq reveals transcriptional dynamics of plant-associated bacteria during root colonization. Nat. Commun. 17, 3021 (2026).

14. Ortega, Á., Zhulin, I. B. & Krell, T. Sensory Repertoire of Bacterial Chemoreceptors. Microbiol. Mol. Biol. Rev. 81, e00033–17 (2017).

15. Sanchis-López, C. et al. Prevalence and Specificity of Chemoreceptor Profiles in Plant-Associated Bacteria. mSystems 6, e00951–21 (2021).

16. Matilla, M. A. & Krell, T. Sensing the environment by bacterial plant pathogens: What do their numerous chemoreceptors recognize? Microb. Biotechnol. 17, e14368 (2024).

17. Matilla, M. A. et al. Chemotaxis of the Human Pathogen Pseudomonas aeruginosa to the Neurotransmitter Acetylcholine. mBio 13, e03458–21 (2022).

18. McKellar, J. L. O., Minnell, J. J. & Gerth, M. L. A high-throughput screen for ligand binding reveals the specificities of three amino acid chemoreceptors from Pseudomonas syringae pv. actinidiae. Mol. Microbiol. 96, 694–707 (2015).

19. Fernandez-López, R., Ruiz, R., de la Cruz, F. & Moncalián, G. Transcription factor-based biosensors enlightened by the analyte. Front. Microbiol. 6, (2015).

20. Bi, S., Pollard, A. M., Yang, Y., Jin, F. & Sourjik, V. Engineering Hybrid Chemotaxis Receptors in Bacteria. ACS Synth. Biol. 5, 989–1001 (2016).

21. Luu, R. A. et al. Hybrid Two-Component Sensors for Identification of Bacterial Chemoreceptor Function. Appl. Environ. Microbiol. 85, e01626–19 (2019).

22. Xu, W. et al. Systematic mapping of chemoreceptor specificities for Pseudomonas aeruginosa. mBio 0, e02099–23 (2023).

23. Xu, W. et al. Specificities of chemosensory receptors in the human gut microbiota. Proc. Natl. Acad. Sci. 122, e2508950122 (2025).

24. Avalos, J. K. et al. Plant pathogenic Ralstonia share two core methyl-accepting chemoreceptors that drive chemotaxis toward distinct amino acid profiles. 2025.04.07.647700 Preprint at 10.1101/2025.04.07.647700 (2025).

25. Matilla, M. A. et al. Is it possible to predict signal molecules that are recognized by bacterial receptors? Environ. Microbiol. 25, 11–16 (2023).

26. Silby, M. W., Winstanley, C., Godfrey, S. A. C., Levy, S. B. & Jackson, R. W. Pseudomonas genomes: diverse and adaptable. FEMS Microbiol. Rev. 35, 652–680 (2011).

27. Sampedro, I., Parales, R. E., Krell, T. & Hill, J. E. Pseudomonas chemotaxis. FEMS Microbiol. Rev. 39, 17–46 (2015).

28. Wiesenbauer, J. et al. Preferential use of organic acids over sugars by soil microbes in simulated root exudation. Soil Biol. Biochem. 109738 (2025) doi:10.1016/j.soilbio.2025.109738.

29. Koh, A., Vadder, F. D., Kovatcheva-Datchary, P. & Bäckhed, F. From Dietary Fiber to Host Physiology: Short-Chain Fatty Acids as Key Bacterial Metabolites. Cell 165, 1332–1345 (2016).

30. Louis, P., Duncan, S. H., Sheridan, P. O., Walker, A. W. & Flint, H. J. Microbial lactate utilisation and the stability of the gut microbiome. Gut Microbiome 3, e3 (2022).

31. Lü, W. et al. The formate channel FocA exports the products of mixed-acid fermentation. Proc. Natl. Acad. Sci. 109, 13254–13259 (2012).

32. Chen, I.-M. A. et al. The IMG/M data management and analysis system v.7: content updates and new features. Nucleic Acids Res. 51, D723–D732 (2023).

33. Wang, G. et al. CRAGE enables rapid activation of biosynthetic gene clusters in undomesticated bacteria. Nat. Microbiol. 4, 2498–2510 (2019).

34. Costa-Gutierrez, S. B. et al. Plant growth promotion by Pseudomonas putida KT2440 under saline stress: role of eptA. Appl. Microbiol. Biotechnol. 104, 4577–4592 (2020).

35. Arslan, E. & Akkaya, Ö. Biotization of Arabidopsis thaliana with Pseudomonas putida and assessment of its positive effect on in vitro growth. Vitro Cell. Dev. Biol. -Plant 56, 184–192 (2020).

36. Nelson, K. E. et al. Complete genome sequence and comparative analysis of the metabolically versatile Pseudomonas putida KT2440. Environ. Microbiol. 4, 799–808 (2002).

37. Nikel, P. I., Chavarría, M., Danchin, A. & de Lorenzo, V. From dirt to industrial applications: Pseudomonas putida as a Synthetic Biology chassis for hosting harsh biochemical reactions. Curr. Opin. Chem. Biol. 34, 20–29 (2016).

38. Volke, D. C., Calero, P. & Nikel, P. I. Pseudomonas putida. Trends Microbiol. 28, 512–513 (2020).

39. Costa-Gutierrez, S. B., Adler, C., Espinosa-Urgel, M. & de Cristóbal, R. E. Pseudomonas putida and its close relatives: mixing and mastering the perfect tune for plants. Appl. Microbiol. Biotechnol. 106, 3351–3367 (2022).

40. Aparicio, T., de Lorenzo, V. & Martínez-García, E. CRISPR/Cas9-enhanced ssDNA recombineering for Pseudomonas putida. Microb. Biotechnol. 12, 1076–1089 (2019).

41. Czajka, J. J. et al. Tuning a high performing multiplexed-CRISPRi Pseudomonas putida strain to further enhance indigoidine production. Metab. Eng. Commun. 15, e00206 (2022).

42. Senthilnathan, R. et al. An update on CRISPR-Cas12 as a versatile tool in genome editing. Mol. Biol. Rep. 50, 2865–2881 (2023).

43. García, V. et al. Identification of a Chemoreceptor for C2 and C3 Carboxylic Acids. Appl. Environ. Microbiol. 81, 5449–5457 (2015).

44. Zhang, X., Hughes, J. G., Subuyuj, G. A., Ditty, J. L. & Parales, R. E. Chemotaxis of Pseudomonas putida F1 to Alcohols Is Mediated by the Carboxylic Acid Receptor McfP. Appl. Environ. Microbiol. 85, e01625–19 (2019).

45. Brewster, J. L. et al. Structural basis for ligand recognition by a Cache chemosensory domain that mediates carboxylate sensing in Pseudomonas syringae. Sci. Rep. 6, 35198 (2016).

46. Upadhyay, A. A., Fleetwood, A. D., Adebali, O., Finn, R. D. & Zhulin, I. B. Cache Domains That are Homologous to, but Different from PAS Domains Comprise the Largest Superfamily of Extracellular Sensors in Prokaryotes. PLOS Comput. Biol. 12, e1004862 (2016).

47. Monteagudo-Cascales, E. et al. Bacterial sensor evolved by decreasing complexity. Proc. Natl. Acad. Sci. 122, e2409881122 (2025).

48. Bainer, R., Park, H. & Cluzel, P. A high-throughput capillary assay for bacterial chemotaxis. J. Microbiol. Methods 55, 315–319 (2003).

49. Berruto, C., Grillo, E., Esturi, S. & Demirer, G. S. A 3D-printed capillary tube holder for high-throughput chemotaxis assays. J. Bacteriol. 208, e00384–25 (2025).

50. Bret, G., Sindt, F. & Rognan, D. Assessing Boltz-2 Performance for the Binding Classification of Docking Hits. J. Chem. Inf. Model. 66, 1511–1521 (2026).

51. Masters, M. R., Mahmoud, A. H. & Lill, M. A. Investigating whether deep learning models for co-folding learn the physics of protein-ligand interactions. Nat. Commun. 16, 8854 (2025).

52. Monteagudo-Cascales, E. et al. Ubiquitous purine sensor modulates diverse signal transduction pathways in bacteria. Nat. Commun. 15, 5867 (2024).

53. Del Valle, I. et al. Translating New Synthetic Biology Advances for Biosensing Into the Earth and Environmental Sciences. Front. Microbiol. 11, (2021).

54. Wang, B. et al. CRAGE-Duet Facilitates Modular Assembly of Biological Systems for Studying Plant– Microbe Interactions. ACS Synth. Biol. 9, 2610–2615 (2020).

55. Berg, H. C. & Turner, L. Chemotaxis of bacteria in glass capillary arrays. Escherichia coli, motility, microchannel plate, and light scattering. Biophysj 58, 919–930 (1990).

56. Abraham, M. J. et al. GROMACS: High performance molecular simulations through multi-level parallelism from laptops to supercomputers. SoftwareX 1–2, 19–25 (2015).

57. Abramson, J. et al. Accurate structure prediction of biomolecular interactions with AlphaFold 3. Nature 630, 493–500 (2024).

58. Eastman, P. et al. OpenMM 8: Molecular Dynamics Simulation with Machine Learning Potentials. J. Phys. Chem. B 128, 109–116 (2024).

59. Huang, J. & MacKerell Jr, A. D. CHARMM36 all-atom additive protein force field: Validation based on comparison to NMR data. J. Comput. Chem. 34, 2135–2145 (2013).

60. Bussi, G., Donadio, D. & Parrinello, M. Canonical sampling through velocity rescaling. J. Chem. Phys. 126, 014101 (2007).

61. Berendsen, H. J. C., Postma, J. P. M., van Gunsteren, W. F., DiNola, A. & Haak, J. R. Molecular dynamics with coupling to an external bath. J. Chem. Phys. 81, 3684–3690 (1984).

62. Essmann, U. et al. A smooth particle mesh Ewald method. J. Chem. Phys. 103, 8577–8593 (1995).

63. Bernetti, M. & Bussi, G. Pressure control using stochastic cell rescaling. J. Chem. Phys. 153, 114107 (2020).

