## Supplementary figures and tables for "A pooled screening approach reveals bacterial chemoreceptors for short-chain carboxylic acids"

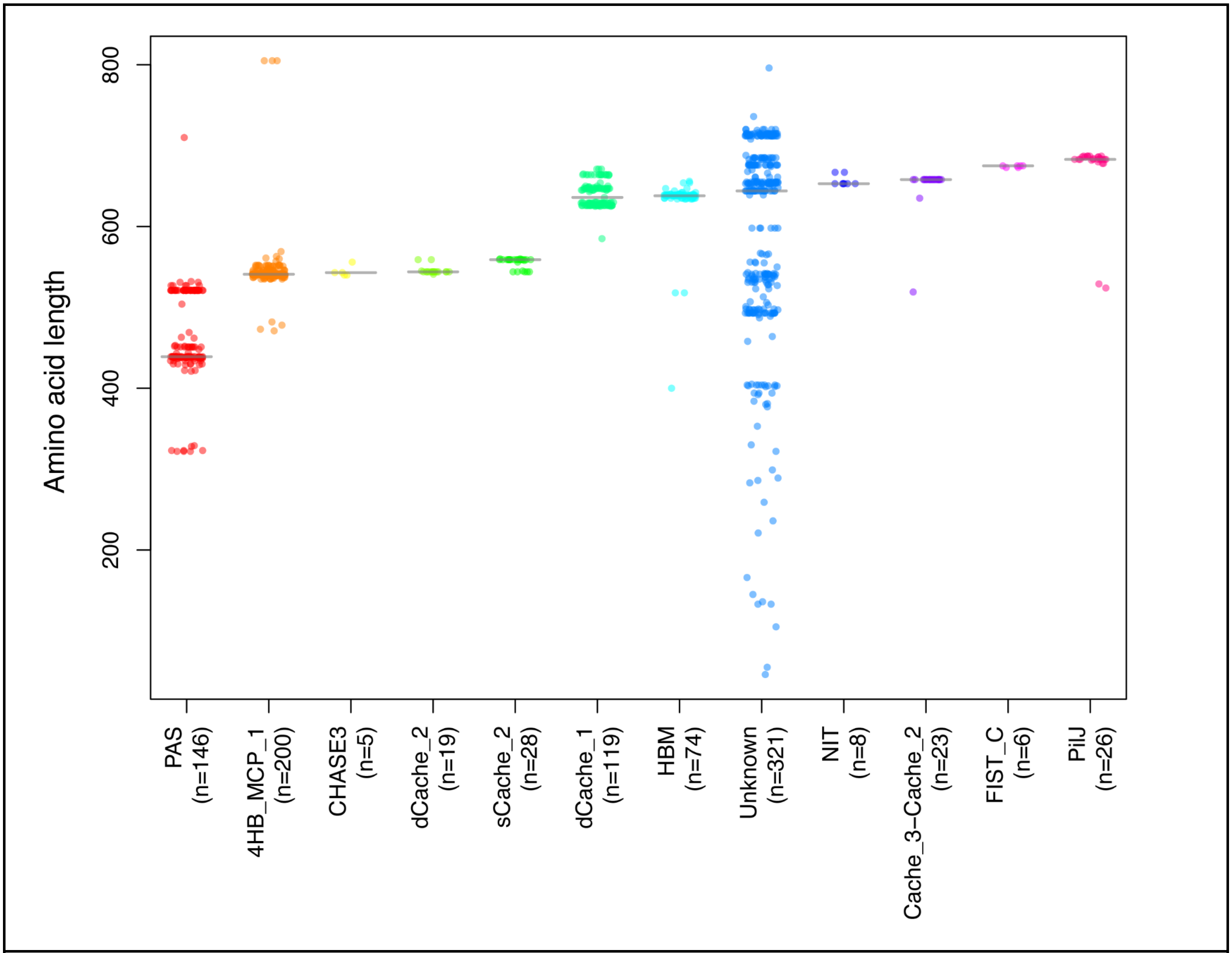

**Figure S1: Structural diversity of LBDs in a *Pseudomonas* chemoreceptor library.**
Individual chemoreceptors are shown as jittered points, colored by LBD classification, with horizontal lines indicating median values for each group. The number of sequences per LBD type is indicated on the x-axis. LBD annotations were assigned based on the JGI-IMG database[1].

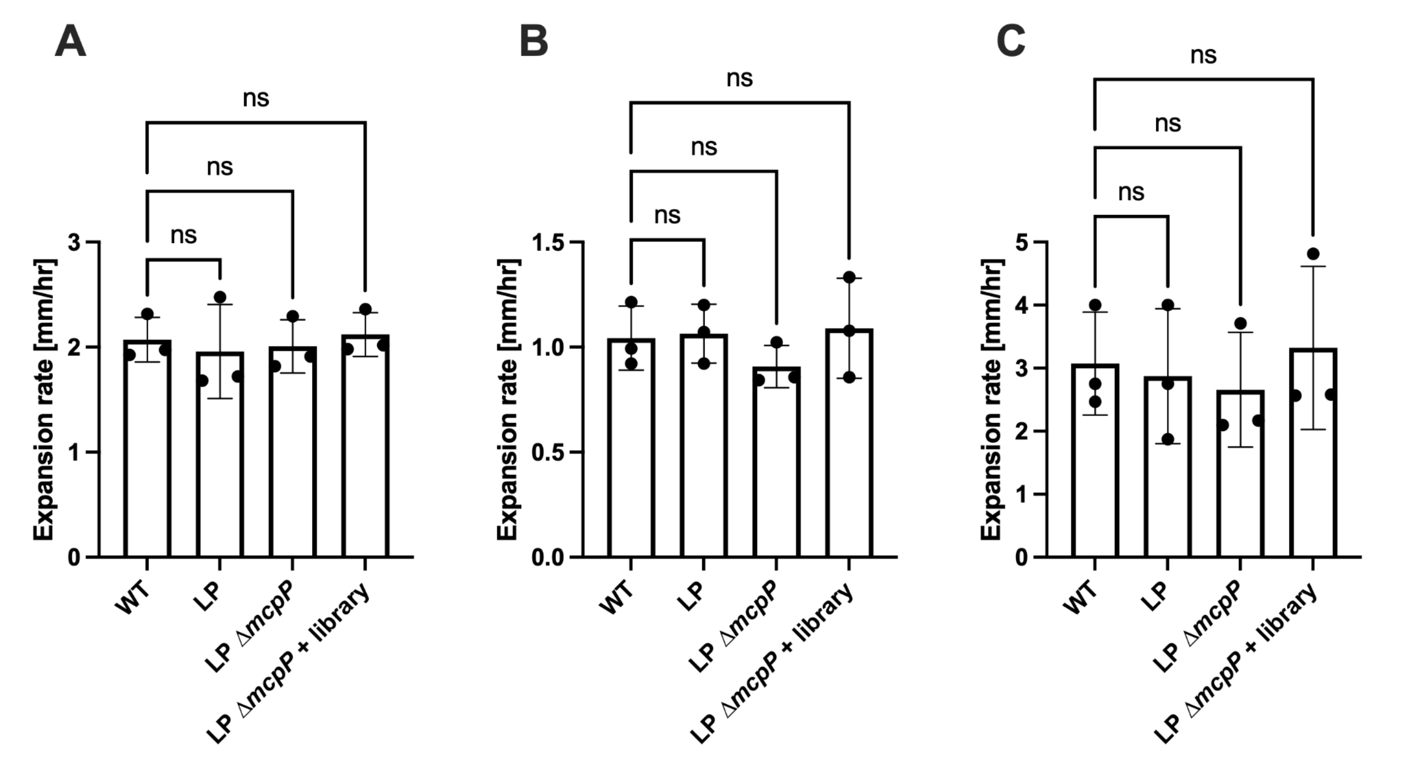

**Figure S2: Comparison of expansion rates between wild-type and engineered strains in soft-agar.** (**A–C**) Expansion rates in soft-agar assays containing 10 mM glucose (A), 10 mM citrate (B), and 2.5% LB (C). Wild type (WT), landing pad integrated strain (LP: strain MC34), Δ*mcpP* mutant (LP Δ*mcpP*: strain MC105), and chemoreceptor library population (LP Δ*mcpP* + library: LS18) were tested (see **Table S2** for a strain list). Data are shown as mean ± s.d. (n = 3 biological replicates). The experiments confirm the genetic engineering process did not alter the motility phenotypes compared to wild-type strain. Statistical significance was assessed using ordinary one-way ANOVA with multiple comparisons; ns, not significant.

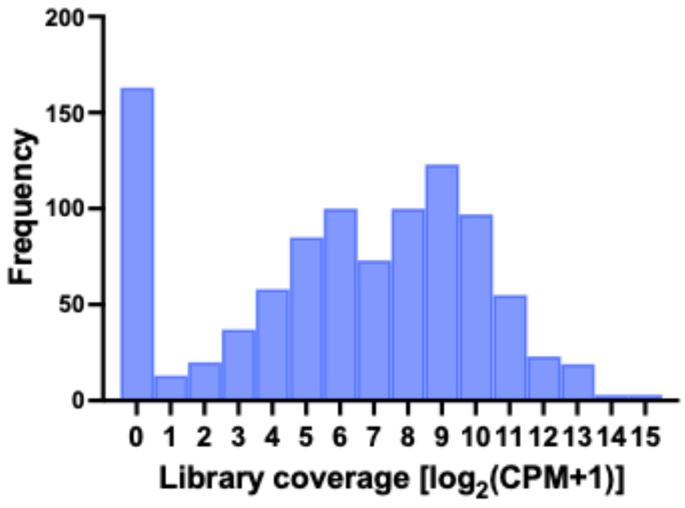

**Figure S3: Coverage of the integrated chemoreceptor gene library.** Amplicon sequencing confirmed successful chromosomal integration of 813 out of 975 chemoreceptor genes in the library population (LS18). Histogram shows the distribution of $\log_2$(mean CPM + 1) values across four replicates. Genes with mean CPM > 0 were considered detected.

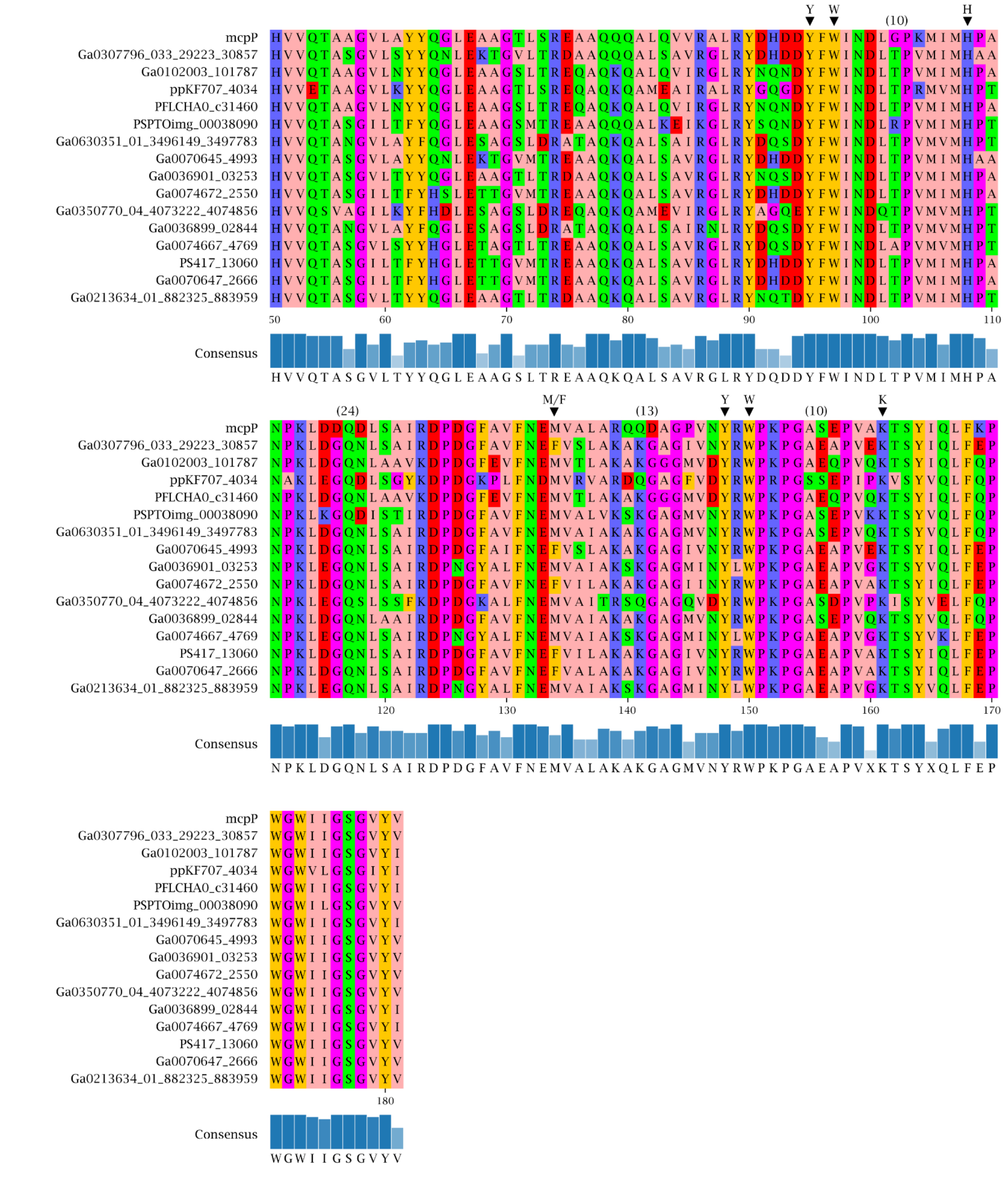

**Figure S4: Multiple sequence alignment of Group I chemoreceptor LBDs with McpP.**
Multiple sequence alignment of ligand-binding domains (LBDs) from McpP (top row) and 15 Group I chemoreceptors showing conservation of key residues implicated in ligand interaction (▼), as identified by Xu *et al*. (2025)[2]. The consensus sequence and residue conservation are shown below the alignment.

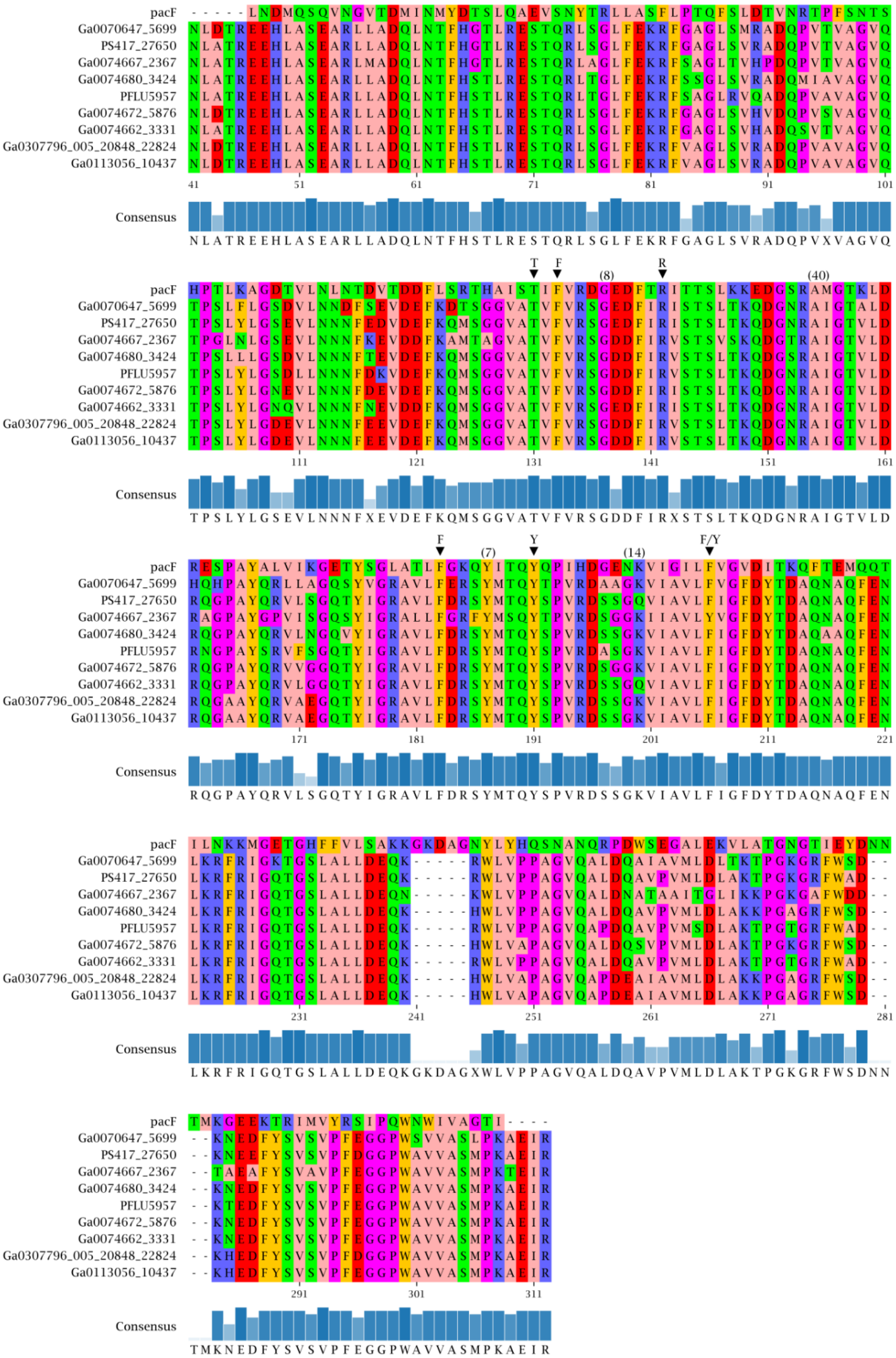

**Figure S5. Multiple sequence alignment of Group II chemoreceptor LBDs with PacF.**
Multiple sequence alignment of ligand-binding domains (LBDs) from PacF (top row) and Group II chemoreceptors showing conservation of key residues implicated in ligand interaction (▼), as identified by Cascales *et al*. (2025)[3]. Despite the low sequence identity of Group II chemoreceptors against PacF, critical residues are conserved across all sequences. The consensus sequence and residue conservation are shown below the alignment.

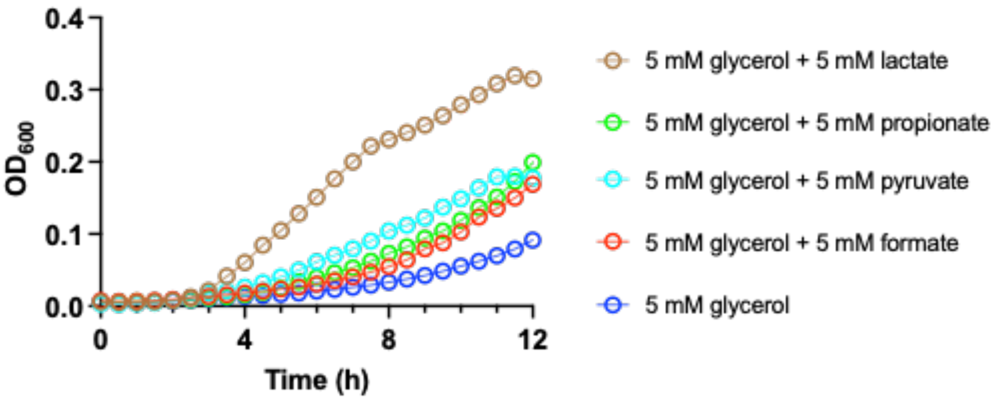

**Figure S6: Growth curves of *P. putida* KT2440 in media supplemented with different carbon sources.** Growth curves in media containing 5 mM glycerol alone or supplemented with 5 mM carboxylic acids. Supplementation with carboxylic acids, including formate, enhanced growth relative to glycerol alone.

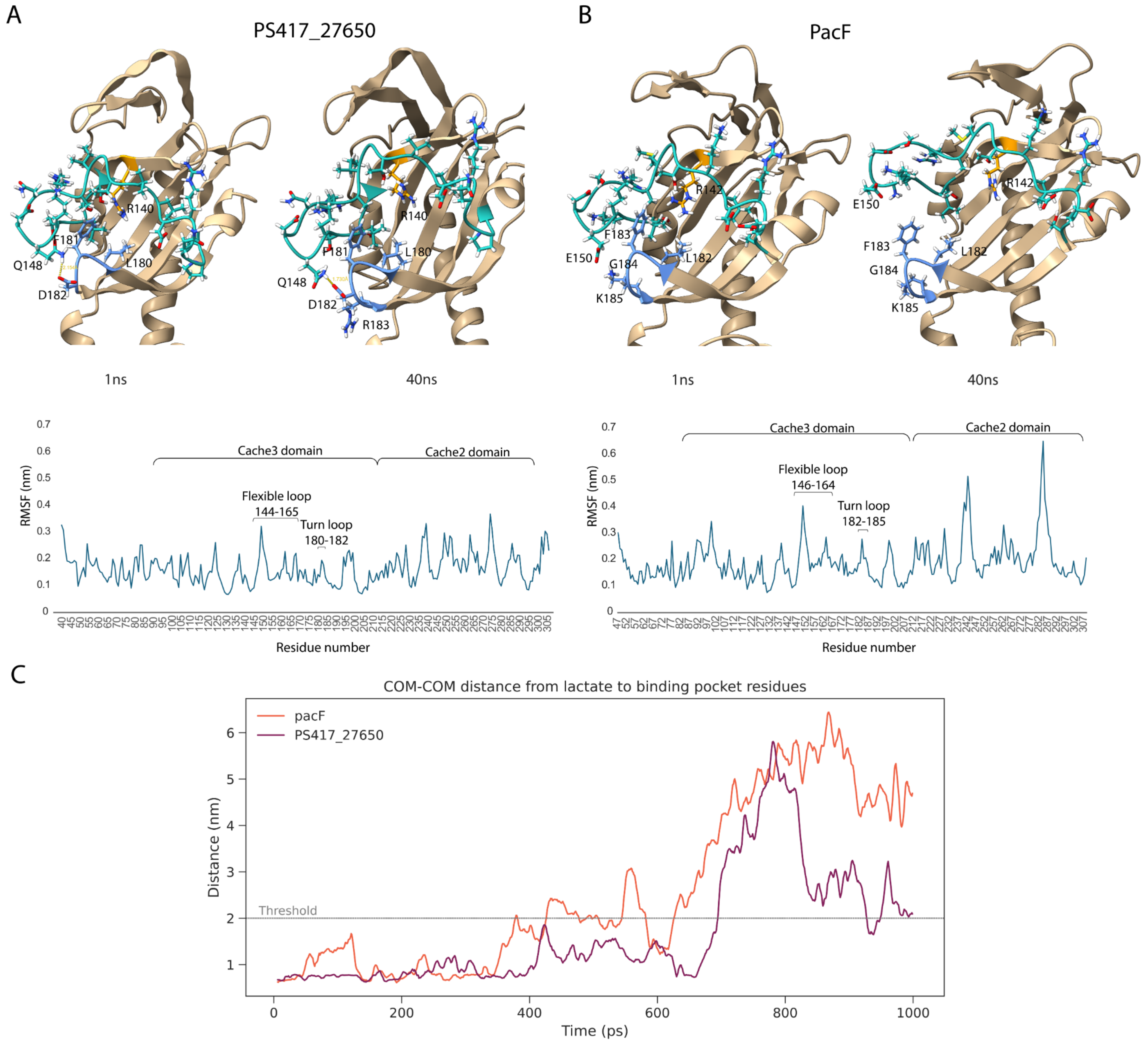

**Figure S7: Molecular dynamics analysis of membrane-distal module of the dCache domain.**
(**A–B**) Representative snapshots at 1 ns and 40 ns showing interactions between residues surrounding the ligand-binding pocket in PS417_27650 (A) and PacF (B). The flexible loop (cyan) and turn loop (blue) are highlighted. The plots below show root mean square fluctuation (RMSF) profiles across residues, indicating differences in loop flexibility. In PS417_27650, Q148 forms interactions with D182 that stabilize the pocket architecture. In contrast, in PacF, the corresponding residues (E150 and G184) do not form stabilizing interactions, and the absence of a side chain at G184 allows increased mobility of F183. This results in displacement of the flexible loop (residues 146–164) relative to the turn loop (residues 182–185), leading to increased tunnel cavity rearrangement over time and reduced stability of lactate within the binding site. (**C**) Consistent with this, molecular dynamics simulations indicate a longer residence time of lactate within the binding pocket of PS417_27650 (6.9 ns) compared to PacF (3.8 ns).

### Supplementary tables

**Table S1: *Pseudomonas* strains used for creating a chemoreceptor gene library.**

| Taxon ID | Isolation site | Strain name | Number of chemoreceptors |
|---|---|---|---|
| 2597489890 | rhizosphere | *Pseudomonas flourescens* Q8-R1 | 46 |
| 8011194704 | rhizosphere | *Pseudomonas fluorescens* 2-79 | 29 |
| 649633086 | rhizosphere | *Pseudomonas fluorescens* SBW25 | 56 |
| 8088461797 | freshwater fish | *Pseudomonas chlororaphis piscium* ATCC 17411 | 29 |
| 2597489887 | rhizosphere | *Pseudomonas chlororaphis aureofaciens* 30-84 | 31 |
| 2585427642 | rhizosphere | *Pseudomonas simiae* WCS417 | 40 |
| 2643221503 | soil | *Pseudomonas* sp. 397MF | 43 |
| 2875505011 | ground water | *Pseudomonas fluorescens* FW300-N2E2 | 45 |
| 2639762715 | drinking water reservoir | *Pseudomonas extremorientalis* LMG 19695 | 43 |
| 2616644927 | phyllosphere | *Pseudomonas lurida* LMG 21995 | 43 |
| 2508501074 | phyllosphere | *Pseudomonas syringae* pv. tomato DC3000 | 49 |
| 2617270901 | pre-filter tanks | *Pseudomonas fluorescens* ATCC 13525 | 42 |
| 2554235341 | rhizosphere | *Pseudomonas protegens* CHA0 | 42 |
| 2713897228 | tomato | *Pseudomonas corrugata* DSM 7228 | 41 |
| 2667528183 | unknown | *Pseudomonas saponiphila* DSM 9751 | 47 |
| 2639762561 | chicory | *Pseudomonas marginalis* NCPPB 667 | 57 |
| 2639762509 | rhizosphere | *Pseudomonas thivervalensis* DSM 13194 | 39 |
| 2639762637 | phyllosphere | *Pseudomonas poae* LMG 21465 | 47 |
| 8110711754 | soil | *Pseudomonas koreensis* DSM 16610 | 33 |
| 8035391570 | cutting fluid | *Pseudomonas oleovorans* NCTC 10692 | 33 |
| 2523533561 | soil | *Pseudomonas taiwanensis* DSM 21245 | 34 |
| 2660238513 | cream | *Pseudomonas synxantha* DSM 18928 | 32 |
| 2531839224 | soil | *Pseudomonas furukawaii* KF707 | 42 |
| 2639762655 | soil | *Pseudomonas kilonensis* DSM 13647 | 39 |

**Table S2: Strains used in this study.**

| Strain ID | Description | Drug resistance |
|---|---|---|
| MC16 | wild type of *P. putida* KT2440 | |
| MC34 | domesticated *P. putida* KT2440 with a landing pad | Km_200 µg/ml |
| MC63 | MC34 with *mcpP* deletion | Km_200 µg/ml |
| MC105 | MC63 with *mVenus* integrated | Apr_200 µg/ml |
| LS18 | MC105 with chemoreceptor library integrated | Apr_200 µg/ml, Tc_10 µg/mL |
| MC111 | MC105 with Ga0074672_5876 | Apr_200 µg/ml, Tc_10 µg/mL |
| MC115 | MC105 with PFLU5957 | Apr_200 µg/ml, Tc_10 µg/mL |
| MC116 | MC105 with Ga0074667_2367 | Apr_200 µg/ml, Tc_10 µg/mL |
| MC117 | MC105 with PS417_27650 | Apr_200 µg/ml, Tc_10 µg/mL |

**Table S3: Plasmids used in this study.**

| Plasmid name | Plasmid details | Marker gene | Description |
|---|---|---|---|
| pEC387 | pR6K-Cre-lox2272-loxP-lox5171 | kanamycin | Constructed in this study |
| pORTMAGE-Pa1 | pTE452-recT-mutL | gentamicin | Addgene (Plasmid #138475) |
| pTE433-beta | pTE433_cpf1 | kanamycin | Gift from Dr. Thomas Eng (Joint BioEnergy Institute) |
| pTE433-mcpP | pTE433_cpf1 with gRNA targeting *mcpP* | apramycin | Constructed in this study |
| pP5171-mVenus | pR6K-loxP-mVenus-lox5171 | apramycin | Constructed in this study |
| pST003 | pR6K-lox2272-loxP | tetracycline | Constructed in this study |

**Table S4: DNA oligos used in this study.**

| Oligo name | Sequence |
|---|---|
| landing_pad_fwd | /5phos/ctgtctcttatacacatctgacacatagatggcgtcgct |
| landing_pad_rev | /5phos/ctgtctcttatacacatctgcgcagaccaaaacgatctc |
| mcpP donor DNA | ggtgaacggcgggggaaattggccggcagggccggccctttcgcgggtgaacccgctcctacg<br>acggcttgtgccgaaccaccgatggacgacgtaggcccatcagctcccgcattgttctttttattgagg<br>tgcatcgaagcatgctttgcagtgagcccacacccgactaatggctaagtgctatgttggtgatagt |
| mcpP_fwd | acatcggcgatccttggaggctacg |
| mcpP_rev | cctggaaggcagtccaaaaatccatcattcc |
| M13-tailed_fwd | /5ammc6/gtaaaacgacggccagtaaggcgactggagtgccatgtccg |
| M13-tailed_rev | /5ammc6/caggaaacagctatgaccgccccactagttctagagcggccg |
| M13_bc1002_F | /5phos/ggtagacacacagactgtgaggtaaaacgacggccagt |
| M13_bc1003_F | /5phos/ggtagacacatctcgtgagaggtaaaacgacggccagt |
| M13_bc1004_F | /5phos/ggtagcacgcacacacgcgcggtaaaacgacggccagt |
| M13_bc1005_F | /5phos/ggtagcactcgactctcgcgtgtaaaacgacggccagt |

| | |
|---|---|
| M13_bc1006_F | /5phos/ggtagcatatatatcagctgtgtaaaacgacggccagt |
| M13_bc1007_F | /5phos/ggtagtctgtatctctatgtggtaaaacgacggccagt |
| M13_bc1008_F | /5phos/ggtagacagtcgagcgctgcggtaaaacgacggccagt |
| M13_bc1009_F | /5phos/ggtagacacacgcgagacagagtaaaacgacggccagt |
| M13_bc1010_F | /5phos/ggtagacgcgctatctcagaggtaaaacgacggccagt |
| M13_bc1011_F | /5phos/ggtagctatacgtatatctatgtaaaacgacggccagt |
| M13_bc1012_F | /5phos/ggtagacactagatcgcgtgtgtaaaacgacggccagt |
| M13_bc1013_F | /5phos/ggtagctctcgcatacgcgaggtaaaacgacggccagt |
| M13_bc1014_F | /5phos/ggtagctcactacgcgcgcgtgtaaaacgacggccagt |
| M13_bc1015_F | /5phos/ggtagcgcatgacacgtgtgtgtaaaacgacggccagt |
| M13_bc1016_F | /5phos/ggtagcatagagagatagtatgtaaaacgacggccagt |
| M13_bc1017_F | /5phos/ggtagcacacgcgcgctatatgtaaaacgacggccagt |
| M13_bc1050_R | /5phos/ggtaggatatacgcgagagagcaggaaacagctatgac |
| M13_bc1051_R | /5phos/ggtagcgtgtctagcgcgcgccaggaaacagctatgac |
| M13_bc1052_R | /5phos/ggtaggtgtgagatatatatccaggaaacagctatgac |
| M13_bc1053_R | /5phos/ggtagctcacgtacgtcacaccaggaaacagctatgac |
| M13_bc1054_R | /5phos/ggtaggcgcacgcactacagacaggaaacagctatgac |
| M13_bc1055_R | /5phos/ggtagcacacgagatctcatccaggaaacagctatgac |
| M13_bc1056_R | /5phos/ggtagagacacacacgcacatcaggaaacagctatgac |
| M13_bc1057_R | /5phos/ggtaggacgagcgtctgagagcaggaaacagctatgac |
| M13_bc1058_R | /5phos/ggtagtgtgtctctgagagtacaggaaacagctatgac |
| M13_bc1059_R | /5phos/ggtagcacacgcactgagatacaggaaacagctatgac |
| M13_bc1060_R | /5phos/ggtaggatgagtatagacacacaggaaacagctatgac |
| M13_bc1061_R | /5phos/ggtaggctgtgtgtgctcgtccaggaaacagctatgac |
| M13_bc1062_R | /5phos/ggtagtctcagatagtctatacaggaaacagctatgac |
| M13_bc1063_R | /5phos/ggtagacacgcatgacacactcaggaaacagctatgac |
| M13_bc1064_R | /5phos/ggtagtatatacagagtcgagcaggaaacagctatgac |
| M13_bc1065_R | /5phos/ggtaggcgctctctcacataccaggaaacagctatgac |

**Table S5. Selected and non-selected chemoreceptors**

| Group | Gene ID | Chemoreceptor origin | Amino acid length | LBD | Screened? |
|---|---|---|---|---|---|
| Group I | 2640002922 | Pseudomonas marginalis NCPPB 667 | 544 | dCache_2 | Yes |
| Group I | 2597858032 | Pseudomonas chlororaphis aureofaciens 30-84 | 544 | dCache_2 | Yes |
| Group I | 2617959847 | Pseudomonas fluorescens ATCC 13525 | 544 | dCache_2 | Yes |
| Group I | 2586031246 | Pseudomonas simiae WCS417 | 544 | dCache_2 | Yes |

| | | | | | |
|---|---|---|---|---|---|
| Group I | 2616887541 | Pseudomonas lurida LMG 21995 | 544 | dCache_2 | Yes |
| Group I | 8088465072 | Pseudomonas chlororaphis piscium ATCC 17411 | 544 | dCache_2 | Yes |
| Group I | 2640374745 | Pseudomonas kilonensis DSM 13647 | 544 | dCache_2 | Yes |
| Group I | 8011198331 | Pseudomonas fluorescens 2-79 | 544 | sCache_2 | Yes |
| Group I | 2532701525 | Pseudomonas furukawaii KF707 | 544 | sCache_2 | Yes |
| Group I | 2597876007 | Pseudomonas fluorescens Q8r1-96 | 544 | dCache_2 | Yes |
| Group I | 8035396207 | Pseudomonas oleovorans NCTC 10692 | 544 | dCache_2 | Yes |
| Group I | 2875505803 | Pseudomonas fluorescens FW300-N2E2 | 544 | dCache_2 | Yes |
| Group I | 2643584364 | Pseudomonas sp. 397MF | 544 | sCache_2 | Yes |
| Group I | 2555670089 | Pseudomonas protegens CHA0 | 545 | sCache_2 | Yes |
| Group I | 2639788514 | Pseudomonas thivervalensis DSM 13194 | 544 | dCache_2 | No |
| Group I | 2662689711 | Pseudomonas synxantha DSM 18928 | 544 | sCache_2 | No |
| Group I | 649636253 | Pseudomonas fluorescens SBW25 | 544 | dCache_2 | No |
| Group I | 2523790463 | Pseudomonas taiwanensis DSM 21245 | 544 | sCache_2 | No |
| Group I | 2716139544 | Pseudomonas corrugata DSM 7228 | 544 | dCache_2 | No |
| Group I | 2640449390 | Pseudomonas extremorientalis LMG 19695 | 544 | dCache_2 | No |
| Group II | 2640452441 | Pseudomonas extremorientalis LMG 19695 | 658 | Cache_3-Cache_2 | Yes |
| Group II | 8011195395 | Pseudomonas fluorescens 2-79 | 658 | Cache_3-Cache_2 | Yes |
| Group II | 2662690122 | Pseudomonas synxantha DSM 18928 | 658 | Cache_3-Cache_2 | Yes |
| Group II | 2640006248 | Pseudomonas marginalis NCPPB 667 | 658 | Cache_3-Cache_2 | Yes |
| Group II | 649639360 | Pseudomonas fluorescens SBW25 | 658 | Cache_3-Cache_2 | Yes |
| Group II | 2640372343 | Pseudomonas kilonensis DSM 13647 | 658 | Cache_3-Cache_2 | Yes |
| Group II | 2640299616 | Pseudomonas poae LMG 21465 | 658 | Cache_3-Cache_2 | Yes |
| Group II | 2616890574 | Pseudomonas lurida LMG 21995 | 658 | Cache_3-Cache_2 | Yes |
| Group II | 2586034105 | Pseudomonas simiae WCS417 | 658 | Cache_3-Cache_2 | Yes |
| Group II | 2671166646 | Pseudomonas saponiphila DSM 9751 | 658 | Cache_3-Cache_2 | No |
| Group II | 2875509614 | Pseudomonas fluorescens FW300-N2E2 | 658 | Cache_3-Cache_2 | No |
| Group II | 2597878614 | Pseudomonas fluorescens Q8r1-96 | 658 | Cache_3-Cache_2 | No |
| Group II | 2617956756 | Pseudomonas fluorescens ATCC 13525 | 658 | Cache_3-Cache_2 | No |
| Group II | 2555672931 | Pseudomonas protegens CHA0 | 658 | Cache_3-Cache_2 | No |
| Group II | 8088468449 | Pseudomonas chlororaphis piscium ATCC 17411 | 658 | Cache_3-Cache_2 | No |
| Group II | 2639792894 | Pseudomonas thivervalensis DSM 13194 | 658 | Cache_3-Cache_2 | No |
| Group II | 2716138758 | Pseudomonas corrugata DSM 7228 | 658 | Cache_3-Cache_2 | No |
| Group II | 2643586187 | Pseudomonas sp. 397MF | 658 | Cache_3-Cache_2 | No |
| Group II | 2597860981 | Pseudomonas chlororaphis aureofaciens 30-84 | 658 | Cache_3-Cache_2 | No |

### References

1. Chen, I.-M. A. *et al.* IMG/M v.5.0: an integrated data management and comparative analysis system for microbial genomes and microbiomes. *Nucleic Acids Res.* **47**, D666–D677 (2019).
2. Xu, W. *et al.* Specificities of chemosensory receptors in the human gut microbiota. *Proc. Natl. Acad. Sci.* **122**, e2508950122 (2025).
3. Monteagudo-Cascales, E. *et al.* Bacterial sensor evolved by decreasing complexity. *Proc. Natl. Acad. Sci.* **122**, e2409881122 (2025).
